# Dentate spikes and external control of hippocampal function

**DOI:** 10.1101/2020.07.20.211615

**Authors:** D. Dvorak, A. Chung, E. H. Park, A. A. Fenton

## Abstract

Mouse hippocampus CA1 place-cell discharge typically encodes current location but during slow gamma dominance (SG_dom_), when slow gamma oscillations (30-50 Hz) dominate mid-frequency gamma oscillations (70-90 Hz) in CA1 local field potentials, CA1 discharge switches to represent distant recollected locations. We report that dentate spike type 2 (DS_M_) events initiated by MECII→DG inputs promote SG_dom_ and change excitation-inhibition coordinated discharge in DG, CA3, and CA1, whereas type 1 (DS_L_) events initiated by LECII→DG inputs do not. Just before SG_dom_, LECII-originating slow gamma oscillations in dentate gyrus and CA3-originating slow gamma oscillations in CA1 phase and frequency synchronize at the DS_M_ peak when discharge within DG and CA3 increases to promote excitation-inhibition cofiring within and across the DG→CA3→CA1 pathway. This optimizes discharge for the 5-10 ms DG-to-CA1 neuro-transmission that coincides with SG_dom_. DS_M_ properties identify extrahippocampal control of SG_dom_, and a cortico-hippocampal mechanism that switches between memory-related modes of information processing.

## Introduction

The hippocampus is critical for long-term memory storage and use, requiring that neural discharge represent both what has occurred and what is happening. How do multifunction neural networks like hippocampus accomplish mutually incompatible tasks like recollecting the past and encoding the present? One possibility is that separate neural circuits operate in parallel to perform each information processing task, but this does not appear to be the case for hippocampus. Rather, in hippocampus the same populations of excitatory and inhibitory neurons are organized such that network discharge patterns, sometimes called cell assemblies (Harris et al., 2003; Hebb, 1949) rapidly switch between different information processing modes, often in a winner-take-all fashion during vicarious-trial-and-error and other choice behaviors (Colgin, 2015; Johnson and Redish, 2007; Kelemen and Fenton, 2010; Kelemen and Fenton, 2013; Kelemen and Fenton, 2016; Papale et al., 2016; Pastalkova et al., 2008; van Dijk and Fenton, 2018; Wu et al., 2017). Indeed, we first reported variability in the discharge of hippocampus place cells that was so extreme it was incompatible with these cells merely signaling the current location within a single cognitive map (Fenton and Muller, 1998; Jackson and Redish, 2007; Olypher et al., 2002a, b) and we went on to show that this variability could be explained as multiple spatial representations during tasks with purposeful behaviors that were directed to specific places (Fenton et al., 2010; Kelemen and Fenton, 2010; Kelemen and Fenton, 2013; Kelemen and Fenton, 2016). In an alternation task, such variability was organized as place representations that alternate within sequences of ∼8 Hz theta oscillations, perhaps reflecting planning between which upcoming alternatives to choose (Kay et al., 2020). We previously reported that position-representing CA1 ensemble spike trains switch between representing the current, local position and distant specific places, which during an active place avoidance task represented recollected locations of prior foot-shock (Dvorak et al., 2018).

Specifically, CA1 discharge switched to signaling distant places during slow-gamma dominance (SG_dom_), when CA1 slow gamma (30-50 Hz) oscillations dominate CA1 mid-frequency (70-90 Hz) gamma oscillations. Now that it is established that such network transitions occur to change hippocampal cognitive information processing, it is essential to understand how such transitions might occur in hippocampus as well as other cognitive networks that transiently switch information processing modes in circumstances that are physically unchanged.

## Results

### Slow gamma dominance in the CA1 LFP switches CA1 place discharge to represent recollection of distant locations

Well-trained mice on the rotating place avoidance arena make evasive movements away from the advancing shock zone, as illustrated in the upper portion of Figure 1A. This behavior demonstrates the mice recollect locations where they were previously shocked. Approximately 1-2 seconds before running away to avoid the location of shock, we observe slow gamma dominance (SG_dom_) in the CA1 LFP, that is the result of a relatively increased rate of slow gamma (30-50 Hz) oscillations and a decreased rate of mid-frequency gamma (70-90 Hz) oscillations (Fig. 1A, lower). The likelihood of SG_dom_ is elevated before mice express active avoidance, with the peak 1.75 s before, when the mouse is often inactive (Fig. 1A, B). In contrast, SG_dom_ is unlikely during the passive approach to the shock zone if the mouse will fail to avoid the approaching shock zone and rather runs away to escape after receiving a shock (Fig. 1B). Such failed avoidances are rare, and most likely occur because the mouse did not recollect the location of shock. Place cells with firing fields in the vicinity of the shock, discharge transiently for about 500 ms during SG_dom_, despite the mouse not being in the vicinity of shock, which can be seen in a single 5-s example (Fig. 1C), in the group data (Fig. 1D), and is confirmed by analysis of place cell overdispersion (Fig. S1A-D). Conversely, place cell ensemble discharge continues to decode to the current location when mice fail to avoid the shock (Dvorak et al., 2018). Because CA1 ensemble discharge can transiently switch from signaling current location to signaling a distant, recollected location, we wondered which network mechanisms can cause this switch between information processing modes (Fig. 1E)?

**Figure 1.**
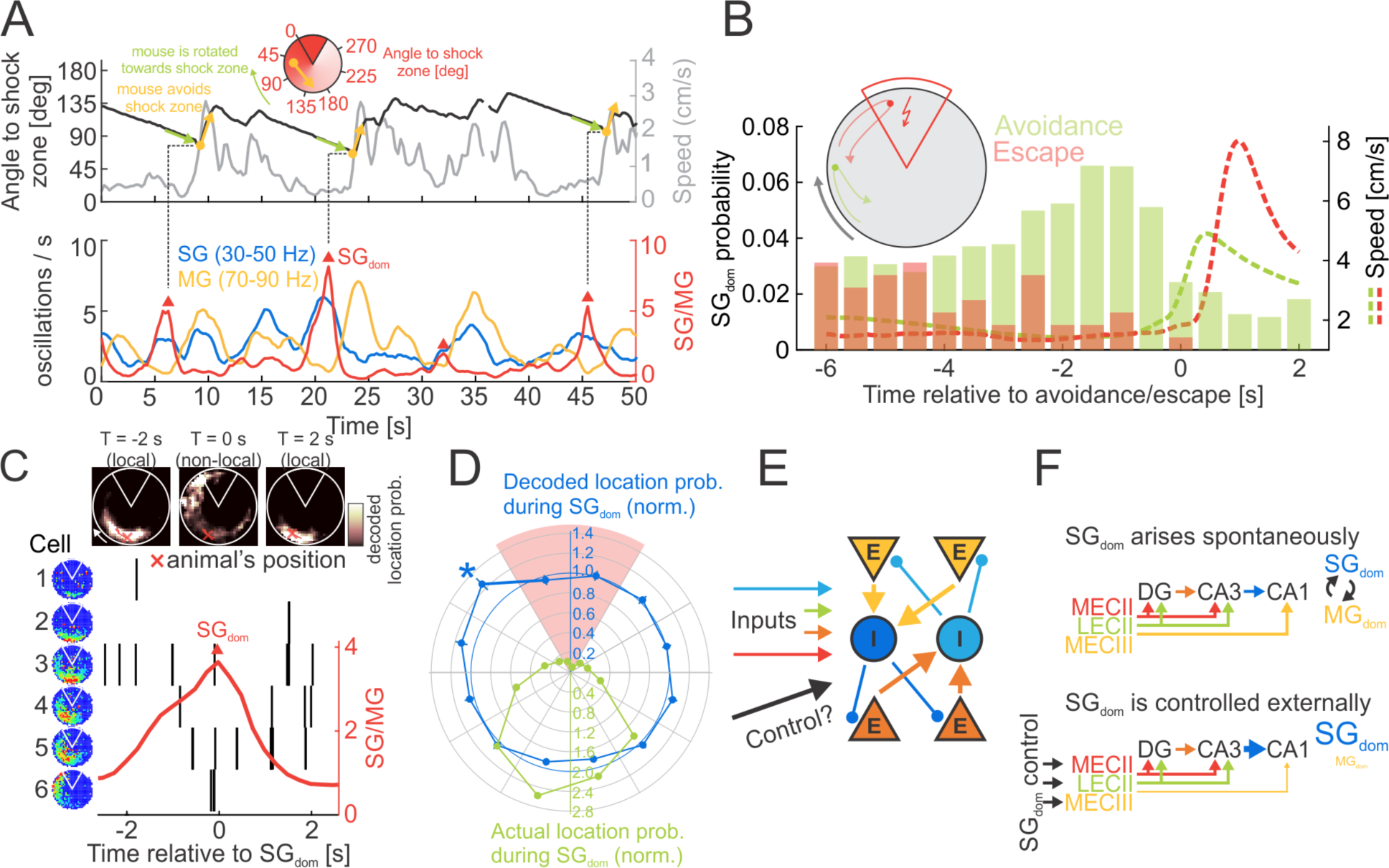
SG_dom_ is a biomarker of memory recollection. A) Avoidances (yellow vectors) mark evasive movements with preceding stillness (green vectors) away from the shock zone without receiving shock. SG_dom_ detected as local maxima (red triangles) in the ratio (red line) of rates of CA1 slow (blue; 30-50 Hz) and mid-frequency (yellow; 70-90 Hz) gamma oscillations, precede avoidance movements by 1-2 seconds. B) SG_dom_ probability histogram before avoidance (green; success = no shock) and escape (red; failure = shocked). C) CA1 single unit discharge (vertical black lines) of a 6-cell ensemble around a SG_dom_ event (red triangle). The firing rate map of each cell is shown on the left. The 2-D posterior probability distributions computed using Bayesian decoding are shown at the top overlayed with the mouse’s 5-s track (red line) and current location (red cross). During SG_dom_, the otherwise accurate Bayesian posterior decodes to the shock zone, away from the mouse’s current location. D) Normalized circular probability distributions of mouse locations (green) and decoded locations (blue) during SG_dom_ (normalization using non-SG_dom_ events). *significant deviation from 1 (t_778_ = 3.10, p = 0.002, Bonferroni’s correction). While SG_dom_ happens predominantly when the mouse is opposite the shock zone (green), discharge during SG_dom_ decodes to locations of shock zone entries (blue). E) Schematic network with winner-take-all dynamics, composed of excitatory (E) and inhibitory (I) neurons, excitatory inputs and a possible external control signal. F) Two hypotheses for hippocampal information processing control (upper) intrinsic, intrahippocampal and (lower) extrinsic, extrahippocampal control. Data in B and D from 2 mice.

One way to switch between multiple mutually-exclusive tasks is to organize the network so that its intrinsic excitation-inhibition dynamics are so balanced that the network spontaneously transitions between multiple information processing modes through intrinsic winner-take-all mechanisms (Fig. 1E, F upper; de Almeida et al., 2009; Rolls and Treves, 1998). In the alternative scenario explored here, SG_dom_-associated switches of the CA1 information processing mode are controlled by discrete events in the perforant path tri-synaptic input from the entorhinal cortex (Fig. 1F, lower).

### Identifying the two types of dentate spike originating from distinct perforant path inputs

CA1 slow gamma originates in CA3 (Lasztoczi and Klausberger, 2014, 2016; Schomburg et al., 2014), motivating us to seek evidence of extrahippocampal control signals in the perforant path projection from ECII to dentate gyrus (DG). We examine dentate spikes (DS), the under-investigated, large amplitude, short duration field potentials that localize to DG (Fig. 2A; Fig. S1E). They result from entorhinal activation; dentate spikes disappear after bilateral removal of entorhinal cortex (Bragin et al., 1995). Similar to sharp-wave ripples (SWR), dentate spikes are synchronized across hemispheres (Bragin et al., 1995; Headley et al., 2017) but in contrast to SWR, DS are thought to cause a synchronized inhibition of granule cells and downstream CA3 and CA1 networks (Fig. 2A left; Penttonen et al., 1997).

**Figure 2.**
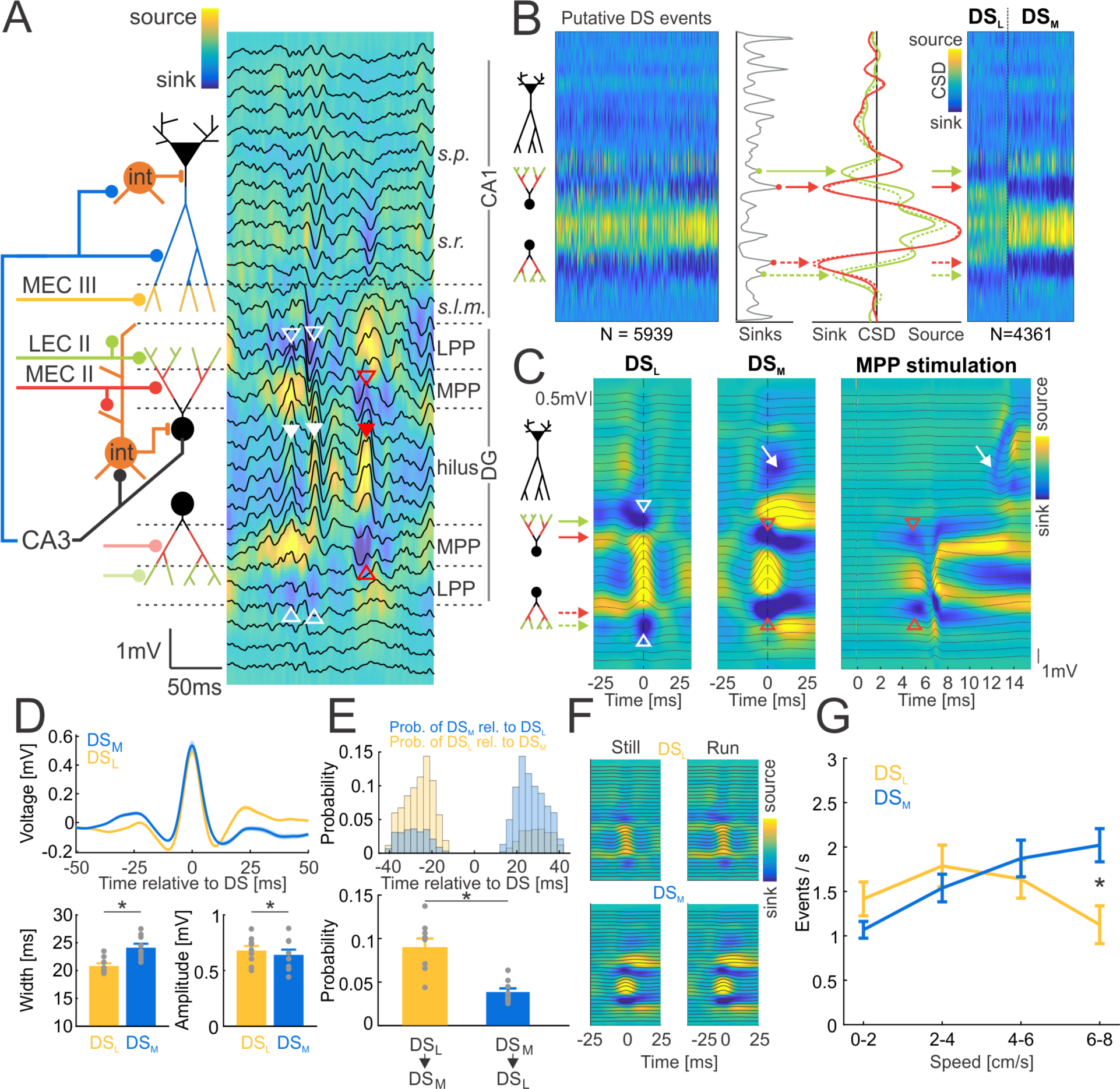
CSD analysis classifies two types of dentate spikes with distinct anatomical, temporal, and behavioral properties. A) DS_L_ (full white arrowheads) is identified by a pair of current-source-density (CSD) sinks in the outer molecular layers of DG (empty white arrowheads) at the LECII projection termination. DS_M_ (full red arrowheads) is identified by a pair of CSD sinks in the middle molecular layers of DG (empty red arrowheads) at the MECII projection termination. Background color represents CSD. Black traces represent the LFP. Schematic (left) illustrating hypothesized components of dentate spike generation and mechanism of CA1 SG_dom_ control. B) CSD of putative DS events (left) and histogram of local minima in CSD profiles (middle) shows peaks aligned with pairs of sink bands in the outer (solid green and dashed green traces) and middle (solid red and dashed red traces) molecular layers. CSD profiles (right) of the two types of DS events. C) Average CSDs of DS_L_ (left) and DS_M_ (middle), with DS_L_ sinks at outer molecular layers (white empty arrowheads) and DS_M_ sinks at middle molecular layers (red empty arrowheads). The CSD of the evoked response to medial perforant path (MPP) stimulation (right), evokes sinks (red empty arrowheads) at the same locations of the DS_M_ sinks. A ∼6 ms latency sink at CA1 *stratum radiatum* occurs after both the MPP evoked response and DS_M_ (white arrow), but not after DS_L_. D) Average LFP of DS_L_ (yellow) and DS_M_ (blue) with their width and amplitude comparisons (bottom). E) Histograms illustrate that DS_M_ often follows DS_L_ (top) and the probability of observing DS_L_→DS_M_ vs. DS_M_→DS_L_ pairs (bottom). F) Average CSDs of DS_L_ (top) and DS_M_ (bottom) during stillness (speed < 2 cm/s; left) and running (speed > 3 cm/s; right). G) Relationship between DS rates and running speed. Averages ± S.E.M. are plotted.

Using current source density (CSD) analysis we classify two types of DS events (Fig. 2B; Star Methods) as DS_L_ (current sink in outer molecular layers of DG) and DS_M_ (current sink in middle molecular layers of DG), corresponding to types 1 and 2 identified in rat (Bragin et al., 1995) and mouse (Buzsaki et al., 2003). Localization of the outer and middle molecular layers of DG, is confirmed by the average CSD of the evoked response to stimulating the medial perforant path (MPP; Fig. 2C, right). The DS_L_ amplitude is larger than DS_M_ (Fig. 2D; paired t_8_ = 2.87, p = 0.02) while DS_M_ width is greater than DS_L_ (paired t_8_ = 8.56, p = 10^-5^). DS_L_ before DS_M_ is more likely than *vice versa* (Fig. 2E; paired t_8_ = 5.61, p = 10^-4^). The average CSDs of DS_L_ and DS_M_ during stillness (speed < 2 cm/s) and running (speed > 3 cm/s) do not visibly differ (Fig. 2F). Rates of DS_L_ and DS_M_ events are not different, but they are differentially modulated by speed (Fig. 2G; 2-way type x speed ANOVA_r.m._, type: F_1,14_ = 0.02, p = 0.88; speed: F_3,12_ = 5.53, p = 0.013; interaction: F_3,12_ = 5.26, p = 0.015, post-hoc tests: DS_M_ > DS_L_ at the greatest speeds of 6-8 cm/s). DS_L_ and DS_M_ are distinct in origin and morphology but not independent, are modulated by behavior, and DS_M_ is more likely to follow DS_L_.

### Dentate spikes modulate oscillatory activity in CA1

We investigate whether DS_L_ and DS_M_ influence the CA1 oscillatory activity components of SG_dom_. Analysis of CA1 oscillatory dynamics using LFP spectral power is confounded by the spectral leakage of DS events in the 30-50 Hz range (Fig. 3A). Accordingly, we use independent component analysis (ICA; Star Methods), which identifies two independent components (IC) in the CA1 LFP below 100 Hz (Fig. 3B), a CA3-originating, *stratum radiatum*-localized slow gamma IC (SG*_SR_* ; mean frequency 34.1 ± 3.0 Hz; Fig. 3C, left; Fig. S2A) and a MECIII-originating, *stratum lacunosum moleculare*-localized mid-frequency gamma IC (MG*_SLM_*; mean frequency 68.9 ± 3.4 Hz; Fig. 3C, right; Fig. S2A).

**Figure 3.**
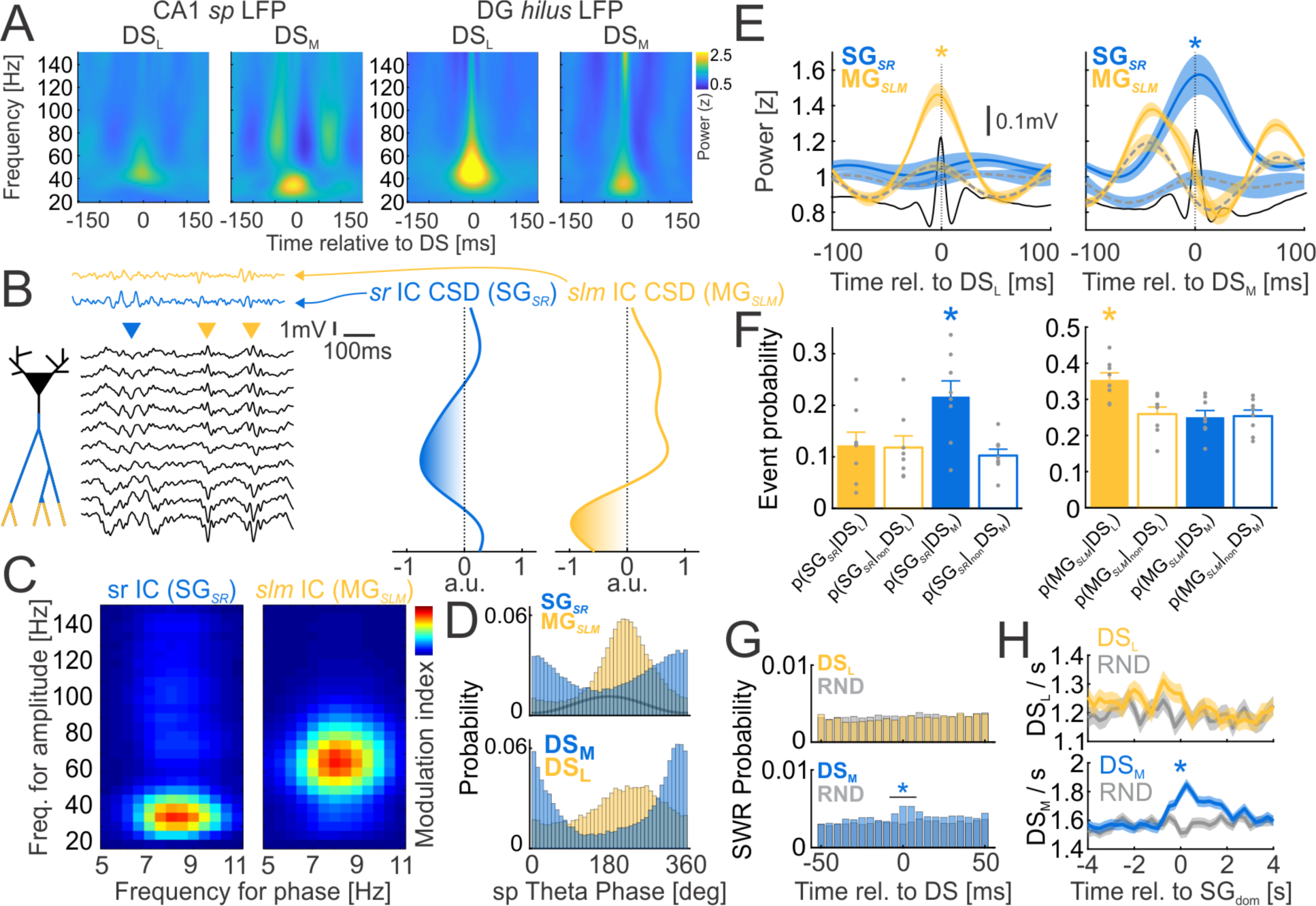
Dentate spikes modulate SG_dom_ and gamma oscillatory activity in CA1. A) Average wavelet spectrogram of CA1 *stratum pyramidale* LFP (left) and DG hilar LFP (right), triggered by DS_L_ and DS_M_ events reveal volume-conducted DS spectral leakage in the 30-50 Hz range, with higher peak frequency associated with narrower DS_L_ events and lower peak frequency associated with wider DS_M_ events, as expected (compare with Fig. 2D). B) ICA decomposition of CA1 LFPs identifies two independent components (IC) < 100 Hz that correspond to *stratum radiatum* dendritic input (SG*_SR_*, CA1 slow gamma, blue), and *stratum lacunosum moleculare* dendritic input (MG*_SLM_*, CA1 mid-frequency gamma, yellow). Arrowheads mark SG*_SR_* (blue) and MG*_SLM_* (yellow) oscillatory events in the LFP. The CSDs of ICA voltage loadings (right) show a SG*_SR_* sink in *stratum radiatum* and a MG*_SLM_* sink in *stratum lacunosum moleculare*. C) Comodulograms between the phase of CA1 *stratum pyramidale* theta (5-11 Hz) and the SG*_SR_* and MG*_SLM_* gamma amplitudes in the 20-150 Hz frequency range show slow gamma peaks for SG*_SR_* and mid-frequency gamma peaks for MG*_SLM_*. D) Theta phase (black line) distribution of SG*_SR_* (blue) and MG*_SLM_* (yellow) oscillatory events (top) compared to theta phase distribution of DS_M_ (blue) and DS_L_ (yellow) events (bottom). E) Power averages of SG*_SR_* (blue) and MG*_SLM_* (yellow) centered on DS_L_ (left) and DS_M_ (right). T=0 marks the DS peak. Gray dashed lines show IC power profiles at random times sampled from the same theta phase distributions as the corresponding DS_L_ and DS_M_ events. Black lines are DS_L_ and DS_M_ event averages. F) Coincidence of the DS and IC oscillatory events detected from SG*_SR_* (left) and MG*_SLM_* (right) at random times. G) SWR probability distribution relative to DS_L_ (top), DS_M_ (bottom) and random times (gray) during stillness. H) Rates of DS_L_ (top) and DS_M_ (bottom) centered at SG_dom_ (color) and randomly selected times (gray). Averages ± S.E.M. are plotted.

If dentate spikes modulate CA1 oscillatory activity, they should systematically co-occur with SG*_SR_* and MG*_SLM_* oscillatory events. SG*_SR_* oscillatory events occur close to the theta trough (339.2° ± 71.4°), whereas the MG*_SLM_* oscillatory events occur close to the theta peak (220.5° ± 59.9°; Fig. 3D top), in agreement with prior work (Fernandez-Ruiz et al., 2017; Lasztoczi and Klausberger, 2014). DS_M_ events occur at the theta trough, coinciding with SG*_SR_* oscillatory events (344.0° ± 59.8; Watson-Williams multi-sample test F_1,15_ = 0.19, p = 0.67), whereas the DS_L_ events occur close to the theta peak, coinciding with the MG*_SLM_* oscillatory events (233.6° ± 71.2°; Watson-Williams multi-sample test F_1,15_ = 3.48, p = 0.08; Fig. 3D).

The theta phase alignment of DS_L_ and MG*_SLM_* events and the distinct phase alignment of DS_M_ and SG*_SR_* events might be expected if DS events control CA1 gamma events, motivating us to determine whether DS events also influence locally-generated CA1 gamma power. We compute DS_L_- and DS_M_-triggered IC power profiles by averaging the z-scored wavelet spectrogram computed from identified ICs across 25-45 Hz for SG*_SR_* and 45-85 Hz for MG*_SLM_* (Fig. 3E). To evaluate if the potential influence of the DS events on the CA1 ICs is distinct from the theta modulation of IC power (see Fig. 3D), we also compare control power profiles triggered by random events that have the same theta phase distribution as the corresponding DS events, but only a chance association with the DS events. Data from 8 of 9 mice are analyzed because one mouse did not have CA1 electrode coverage. MG*_SLM_* is increased 36% at the DS_L_ peak compared to random (paired t_7_ = 6.82, p = 10^-4^), while SG*_SR_* is not (paired t_7_ = 0.35, p = 0.7; Fig. 3E, left). In contrast, SG*_SR_* is increased 67% at the peak of DS_M_ compared to random (paired t_7_ = 5.0, p = 0.002), while MG*_SLM_* is decreased during DS_M_ (paired t_7_ = 2.68, p = 0.03; Fig. 3E, right).

To examine the co-occurrence of the DS and CA1 gamma oscillatory events, we randomly pick 1000 times from each 30-min recording and assess whether MG*_SLM_* or SG*_SR_* occurs within a 50-ms coincidence interval of DS_L_ or DS_M_. The probability of observing SG*_SR_* is greater if DS_M_ is observed [Fig. 3F, left; F_3,31_ = 9.59, p = 10^-4^, p(SG*_SR_* | DS_M_) > p(SG*_SR_* | DS_L_) = p(SG*_SR_* | nonDS_M_) = p(SG*_SR_* | nonDS_L_)] while the probability of observing MG*_SLM_* is greater if a DS_L_ event is observed [Fig. 3F, right; F_3,31_ = 7.31, p = 10^-4^, p(MG*_SLM_* | DS_L_) > p(MG*_SLM_* | DS_M_) = p(MG*_SLM_* | nonDS_L_) = p(MG*_SLM_* | nonDS_M_)].

Since DS_M_ increases the power of CA3-originating SG*_SR_* (Fig. 3E) and dentate spikes co-occur with CA3-originating sharp wave ripples (SWR; Bragin et al., 1995), we computed the probability of a SWR within ±50 ms of DS_L_, DS_M_, and random events during stillness (speed < 2 cm/s; Fig. 3G). SWR probability is increased ±10 ms of the DS_M_ peak but not DS_L_ (Fig. 3G; One sample test for proportions: p(SWR | DS_L_) = 0.012, z = 0.71, p = 0.3; p(SWR | DS_M_) = 0.018, z = 13.20, p = 10^-39^).

If DS_M_ controls CA1 information processing mode, these findings of DS modulation of CA1 gamma predict that DS_M_ (but not DS_L_) promotes CA1 SG_dom_. We evaluated this prediction using SG_dom_ events collected during place avoidance behavior, where SG_dom_ events identify recollection (Fig. 1). The rate of DS_M_ but not DS_L_ events is elevated at the time of SG_dom_ (Fig. 3H; DS_L_: t_5255_ = 1.81, p = 0.07; DS_M_: t_5255_ = 5.07, p < 0.0001).

### Dentate spikes modulate individual cycles of CA1 gamma oscillations, DS_M_ promoting SG_dom_

If DS_M_ events control CA1 information processing by promoting SG_dom_, then DS_M_ should influence CA1 gamma oscillations with a precision comparable to the ∼6 ms conduction time from dentate gyrus to CA1 (Fig. 2C, white arrows). Because measuring gamma power requires ∼100 ms (3-5 cycles of an oscillatory burst; Figs. 3E, S2C), and spiking is most likely during oscillatory minima (Fig. S2B; Dvorak and Fenton, 2014; Lasztoczi and Klausberger, 2016; Schomburg et al., 2012), we measure discrete oscillatory events with ∼15 ms resolution, as the local minima of oscillatory bursts (Fig. 4A inset; Star Methods). The findings in Fig. 4, data acquired during the place avoidance task, are essentially similar in home-cage data (Fig. S3A).

**Figure 4.**
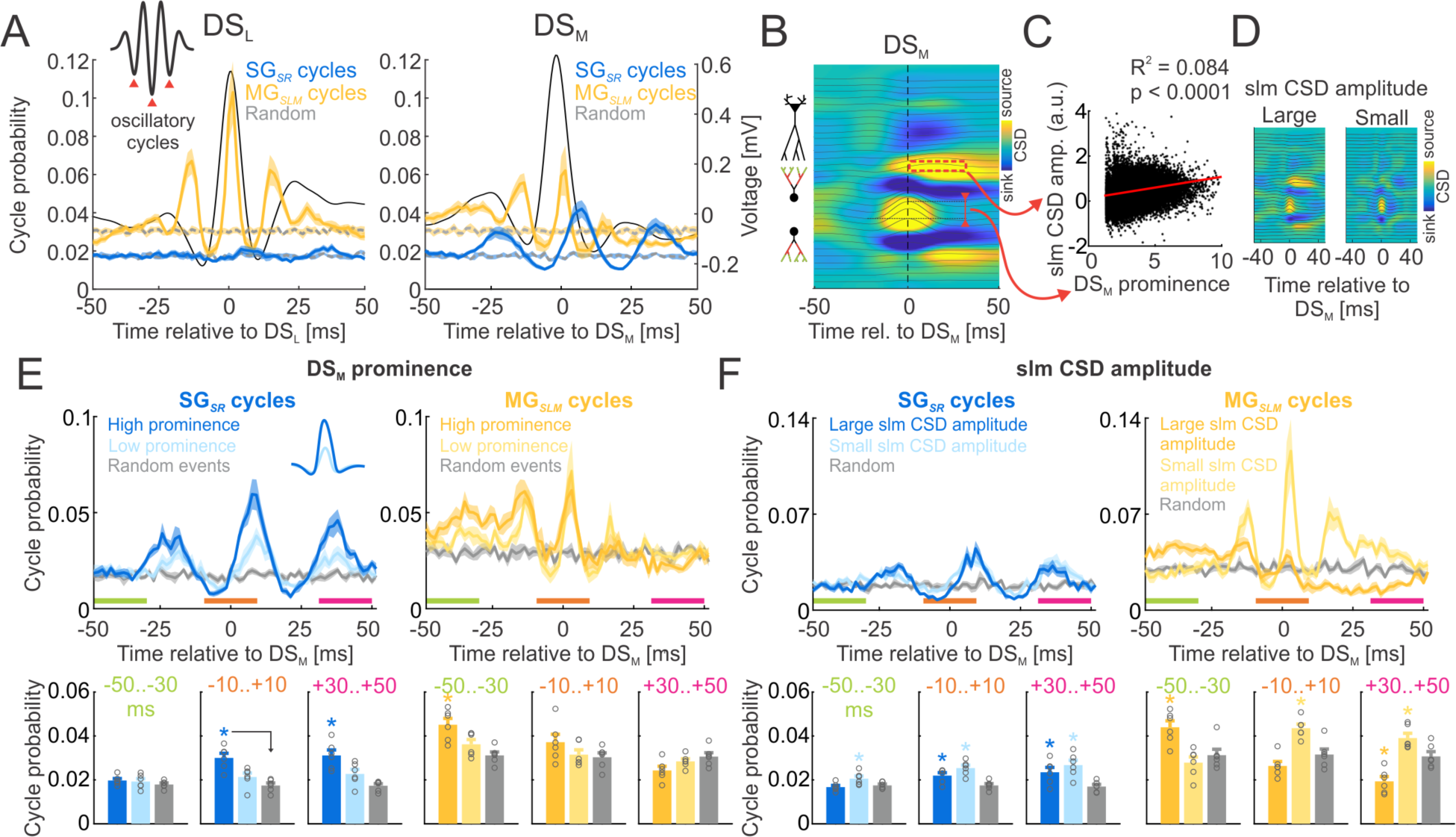
DS_M_ controls the oscillatory components of SG dominance. A) Probability of oscillatory cycles (inset, red arrowheads) detected in SG*_SR_* (blue) and MG*_SLM_* (yellow) relative to DS_L_ (left), DS_M_ (right) and random events (gray). Average DS waveforms are black. B) CSD indicating DS_M_ prominence (red line with reversed arrows) and amplitude of the *slm* CSD source that follows DS_M_ (red rectangle). C) Scatter plot of DS_M_ prominence versus the *slm* CSD source during DS_M_, with linear fit (red). D) Average DS_M_ CSD of the 10% largest (left) and smallest (right) *slm* CSD sources; the DS_M_ prominence is similar. E) Probability of SG*_SR_* cycles (blue; left) and MG*_SLM_* cycles (yellow; right) during the 10% largest (dark color), and smallest (light color) DS_M_ (E) prominence, (F) *slm* CSD source amplitude; random events (gray) and comparisons BEFORE (-50 to -30 ms), DURING (-10 to +10 ms) and AFTER (30 to 50 ms) the DS_M_ events. Averages ± S.E.M. are plotted.

The probability of observing a MG*_SLM_* oscillatory cycle is strongly enhanced ±10 ms of the DS_L_ peak (paired t_7_ = 5.43, p = 0.001; Fig 4A, left) as well as 15 ms before and 16 ms after the DS_L_ peak, corresponding to a MG*_SLM_* oscillatory frequency of 69 Hz, whereas the probability of observing a SG*_SR_* oscillatory cycle remains unchanged during DS_L_ (paired t_7_ = 0.69, p = 0.52; Fig 4A, left). In contrast, the probability of observing a SG*_SR_* oscillatory cycle is enhanced 6 ms after the peak of DS_M_, corresponding to the transmission time between DG and CA1, which is primarily influenced by the CA3→CA1 synaptic delay (Mizuseki et al., 2012; Fig 4A, right; paired t_7_ = 4.52, p = 0.003). The probability of a SG*_SR_* oscillatory cycle is also enhanced 24 ms before and 33 ms after the DS_M_ peak corresponding to a SG*_SR_* oscillatory frequency of 34 Hz, whereas the probability of observing a MG*_SLM_* oscillatory cycle is not different (Fig. 4A right; paired t_7_ = 0.79, p = 0.45;). There is a non-oscillatory increase of the MG*_SLM_* oscillatory cycle probability 30-50 ms before DS_M_ (Fig 4A, right; paired t_7_ = 3.80, p = 0.007) and a reduced probability of observing a MG*_SLM_* oscillatory cycle 30-50 ms after DS_M_ (Fig 4A, right; paired t_7_ = 7.39, p = 10^-4^).

The consequences of MPP manipulations are not straightforward (Brun et al., 2002; Garner et al., 2012; Kanter et al., 2017; Miao et al., 2015; Schlesiger et al., 2018), which was confirmed by chemogenetic silencing, electrical stimulation of MPP, and anesthesia (Fig. S4). Consequently, to rigorously test the hypothesis that DS_M_ promotes SG_dom_, we examine whether spontaneously strong and weak DS_M_ events differentially promote SG_dom_. Because DS_M_ can both increase the likelihood of SG*_SR_* and attenuate the likelihood of MG*_SLM_* to promote SG_dom_, DS_M_ were classified according to their prominence (Fig. 4B) and independently, by the ∼10ms post-DS_M_ CSD source that suggests increased inhibition, corresponds to the DG→CA1 transmission time, and localizes to the vicinity of the hippocampal fissure and CA1 *slm* (red rectangle in Fig. 4B).

Because the *slm* CSD source accounts for only 8% of the variance in DS_M_ prominence (Fig. 4C, D; r_2_ = 0.084, p < 0.0001), we used both features to evaluate the causal predictions that 1) DS_M_ events with a large prominence will increase SG*_SR_* and 2) that DS_M_ events with a large *slm* CSD source will decrease MG*_SLM_*, each promoting SG_dom_.

The probability of observing SG*_SR_* and MG*_SLM_* oscillatory cycles was computed in relation to the 10% of DS_M_ with the highest and lowest prominence, DURING ±10 ms of the DS_M_ peak (orange bar in Fig. 4E), 30-50 ms BEFORE (green bar in Fig. 4E), and 30-50 ms AFTER (magenta bar in Fig. 4E). CA1 SG*_SR_* oscillatory cycles were more likely DURING (F_2,17_ = 13.10, p = 0.0005, high > low = random) and AFTER (F_2,17_ = 12.15, p = 0.0007, high > low = random) the high prominence DS_M_ events compared to the low prominence DS_M_ and random events. These patterns were not observed in relation to DS_L_ events (Fig. S3C). CA1 SG*_SR_* oscillatory cycles were most probable ∼10 ms after the high and low prominence DS_M_ peaks, similar to the DG →CA1 transmission time. In contrast, CA1 MG*_SLM_* oscillatory cycles were more likely BEFORE high prominence DS_M_ (F_2,17_ = 8.32, p = 0.004, high > low = random) but not AFTER (F_2,17_ = 2.73, p = 0.09). Both the high and low prominence DS_L_ events increased the probability of MG*_SLM_* oscillatory cycles during DS_L_ (Fig. S3D). Together these findings further support the hypothesis that DS_M_ controls SG_SR_ to promote SG_dom_ in CA1.

Complementary patterns of promoting SG_dom_ are evident when DS_M_ events are categorized as being the 10% with the largest or smallest *slm* CSD source (Fig. 4F). CA1 MG*_SLM_* oscillatory cycles were more likely BEFORE (F_2,17_ = 9.57, p = 0.002, large > small = random), and less likely DURING (F_2,17_ = 16.93, p = 0.0001, small > large = random) and AFTER (F_2,17_ = 20.68, p < 0.0001, small > random > large) DS_M_ events with large *slm* CSD source (Fig. 4F right). CA1 SG*_SR_* oscillatory cycles were less likely BEFORE (F_2,17_ = 5.39, p = 0.017, small > large = random), DURING (F_2,17_ = 11.42, p = 0.001, small > large > random) and AFTER (F_2,17_ = 7.00, p = 0.0071, small > large > random) DS_M_ events with large *slm* CSD source (Fig. 4F left). Taken together, these analyses confirm the causal predictions that the prominence of DS_M_ and the amplitude of the associated *slm* CSD source together control SG*_SR_* and MG*_SLM_* gamma oscillations to promote SG_dom_.

### DS_M_ synchronizes DG and CA1 slow gamma band oscillations

Dentate DS_M_ events increase CA3-originating SG*_SR_* to promote SG_dom_, but is CA3 activity under enhanced or reduced DG influence during DS_M_? We start by studying the synchrony of DG and CA1 oscillations during DS_L_ and DS_M_. ICA combined with CSD-based classification of DS events disentangles DS events and DG oscillatory components that both originate in the perforant path projection to DG (Fig. S2D-M; Barth et al., 2018; Fernandez-Ruiz et al., 2013).

ICA identified a lateral perforant path (LPP) IC localized to the outer molecular layer DG sinks in the CSD (Fig. S2M) of the ICA voltage loadings and has a slow gamma peak in the CA1 theta phase comodulogram (SG_LPP_; Fig. 5A, bottom left; mean frequency 43.9 ± 5.0 Hz). The medial perforant path (MPP) IC (Fig. 5A, right) localized to the middle molecular layer DG sinks in the CSD (Fig. S2M) of the ICA voltage loadings and has a mid-frequency gamma peak in the CA1 theta phase comodulogram (MG_MPP_; Fig. 5A, bottom right; mean frequency 71.0 ± 2.7 Hz).

**Figure 5.**
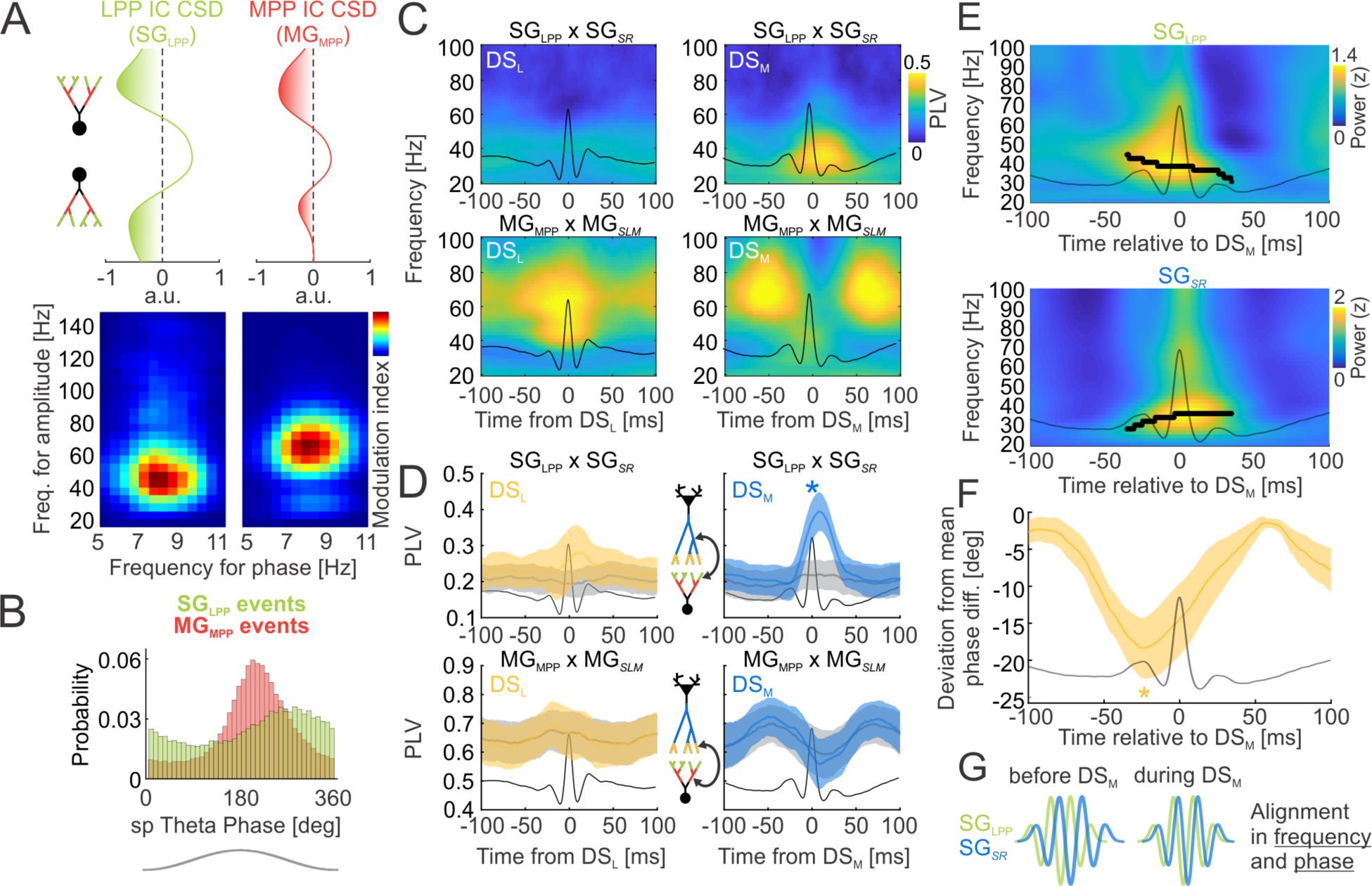
DS_M_ synchronizes the slow gamma oscillatory inputs from lateral perforant path to dentate gyrus and from CA3 to CA1 *stratum radiatum*. A) CSDs of ICA voltage loadings (top) of the LPP IC (SG_LPP_; green), and the MPP IC (MG_MPP_; red) in the DG. Comodulograms (bottom) between the phase of CA1 theta (5-11 Hz) and the amplitude of both IC components across 20-150 Hz. B) Theta phase distribution of SG_LPP_ (green) and MG_MPP_ (red) oscillatory events. C) Example of phase locking value (PLV) between DG and CA1 slow gammas (SG_LPP_ x SG*_SR_*; top) and between DG and CA1 mid-frequency gammas (MG_MPP_ x MG*_SLM_*; bottom) time-locked to DS_L_ (left) and DS_M_ (right). D) Group PLV measures averaged across slow gamma (25-45 Hz for SG_LPP_ and SG*_SR_*) and mid-frequency gamma (45-85 Hz for MG_LPP_ and MG*_SLM_*). Gray: PLV profiles of random samples from the corresponding DS_L_ and DS_M_ theta phase distributions. E) Average wavelet spectrogram of SG_LPP_ (top) and SG*_SR_* (bottom) around the time of DS_M_ (T = 0 ms). Black dots indicate frequency of peak power at each time point ±40 ms. F) Group average of how much the instantaneous phase differences between SG_LPP_ and SG*_SR_* differ from the mean phase difference. Averages ± S.E.M. and average DS waveforms (black) are plotted. G) Schematic of frequency and phase alignment of SG_LPP_ and SG*_SR_* during DS_M_.

While the mean frequency of dentate SG_LPP_ is higher than the mean frequency of the CA1 SG*_SR_* (paired t_6_ = 3.69, p = 0.01), the mean frequencies of dentate MG_MPP_ and CA1 MG*_SLM_* do not differ (paired t_6_ = 2.17, p = 0.07).

CA1 theta is used as an intrinsic network time reference to analyze the phase preference of the dentate SG_LPP_ and MG_MPP_ oscillations (Fig. 5B). Dentate SG_LPP_ oscillations occur at a late descending phase, close to the theta trough (277.4° ± 74.3°) that precedes both DS_M_ (Watson-Williams multi-sample test F_1,14_ = 6.3, p = 0.03) and SG*_SR_* (Watson-Williams multi-sample test F_1,14_ = 4.5, p = 0.05). Dentate MG_MPP_ occurs close to the theta peak (214.9° ± 59.8°), similar to DS_L_ (Watson-Williams multi-sample test F_1,14_ = 1.9, p = 0.19) as well as CA1 MG*_SLM_* oscillations (Watson-Williams multi-sample test F_1,14_ = 0.04, p = 0.83).

Because the DG SG_LPP_ and the CA1 SG*_SR_* oscillations appear at similar phases of the CA1 theta cycle, and the DG MG_MPP_ and CA1 MG*_SLM_* also appear at similar phases of CA1 theta (Figs. 3D, 5B), dentate spikes could synchronize the DG and CA1 subfields. We test this possibility by measuring the phase coupling between DG and CA1 slow and mid-frequency gamma oscillations. The frequency-band specific phase locking values (PLV; Lachaux et al., 1999) time-locked to DS events (Fig. 5C; Star Methods) show that the SG_LPP_ and SG*_SR_* oscillations are not coupled during DS_L_ (Fig. 5C, top left), whereas MG_MPP_ and MG*_SLM_* couple during DS_L_ (Fig. 5C, bottom left). In contrast, the SG_LPP_ and SG*_SR_* couple strongly after the DS_M_ peak (Fig. 5C, top right), and MG_MPP_ and MG*_SLM_* couple ∼50 ms before and ∼75 ms after DS_M_ (Fig. 5C, bottom right). The phase interactions are frequency-specific, especially in the case of the DS_M_-triggered events. Consequently, PLV was averaged across slow 25-45 Hz and mid-frequency 45-85 Hz gamma bands (Fig. 5D), and to evaluate whether or not any DS-related phase coupling between the DG and CA1 gamma oscillations is simply a result of the co-occurrence of DS events and gamma oscillations at similar theta phases (Figs. 3D, 5B), we also compute PLV around randomly selected time points that are sampled from the theta phase distributions of the DS_L_ and DS_M_ events (gray profiles in Fig. 5D). The only significant departure from random was during DS_M_, between the SG_LPP_ and the SG*_SR_* oscillations (Fig. 5D, top right; paired t_6_ = 4.04, p = 0.006). The peak of this phase locking occurs 9 ms after the DS_M_ peak. Similarly, the DS_M_-locked SG*_SR_* oscillatory cycles lag the SG_LPP_ oscillatory cycles by 6 ms (Fig. S3A, B) pointing again to the DG→CA1 transmission time that was observed in Figs. 2C and 4A.

### CA1 SG_SR_ and DG SG_LPP_ are frequency- and phase-tuned during DS_M_

Given a fixed duration of the gamma-generating GABA_A_ receptor response, the frequency of a gamma oscillation can be adjusted by changing the level of network excitation, such that greater excitation produces faster oscillations because GABA inhibition can be overcome sooner (Whittington et al., 1995). Because the 44-Hz SG_LPP_ and the 34-Hz SG*_SR_* oscillate at different frequencies (Figs. 3C, 5A), but phase lock during DS_M_ (Fig. 5D), the gamma-generating mechanism predicts input-driven changes in both the frequency and phase relationships for the phase alignment during DS_M_. We analyze the frequency relationships of SG_LPP_ and SG*_SR_* during DS_M_ to test the predictions, During DS_M_, the frequency of SG_LPP_ decreases from 43 Hz to 36 Hz at the peak of DS_M_, whereas the frequency of SG*_SR_* increases from 28 Hz to 36 Hz at the peak of DS_M_, effectively aligning the frequencies of the DG and CA1 originating oscillations (Fig. 5E). Analysis of the phase relationships of SG_LPP_ and SG*_SR_* during DS_M_ (Fig. 5F) shows the maximum deviation from the mean phase difference occurs 25 ms before the DS_M_ peak (t_6_= 4.51, p = 0.004), and the phase offset reverts to the mean phase difference by 60 ms after the DS_M_ peak (t_6_= 2.37, p = 0.06). At the peak of DS_M_, the phase difference is reduced by 14.5±12.8° (from 11 to 9 ms), similar to the DG→CA1 transmission time observed in Figs. 2C, 4A, 5D and S3A.

### DS_M_ increases DG, CA3 and CA1 discharge rates and cofiring

The hypothesis that DS_M_ has a causal role in promoting SG_dom_ (Figs. 3, 4), and synchronizing slow gamma oscillations at the LPP terminals in DG and CA3 terminals in CA1 (Fig. 5), predicts that DS_M_ organizes DG, CA3, and CA1 discharge. Objectively classified, presumptive principal cells (E) and narrow waveform interneurons (In) were localized and studied to test the prediction (Fig. 6A, B; Star Methods). We compute the firing rates of presumptive granule cells (GC, n = 141), mossy cells (MC, n = 140), CA3 (n = 104) and CA1 (n = 145) principal cells as well as narrow waveform interneurons detected in their proximity (n = 435) during 10 ms windows shifted relative to DS_L_ and DS_M_ events (Fig. 6C). These are compared with the firing rates at random times. DS events contaminated by SWR events were excluded to minimize potential SWR bias (Fig. 3G). During DS_L_ the discharge of GC decreases by 13% (t_140_ = 2.92, p = 0.004). Similarly, the discharge of MC decreases by 19% (t_139_ = 3.54, p = 0.0005). CA3 and CA1 principal cells did not change firing rates (Fig. 6C; CA3: t_103_ = 1.36, p = 0.18; CA1: t_144_ = 1.49, p = 0.14). In contrast, during DS_M_, GC rates increase by 106%, MC rates increase by 117%, and CA3 rates increase by 47%, whereas CA1 principal cell rates do not significantly increase as observed during SG_dom_ (Fig. 6C; GC: t_140_ = 5.82, p = 10^-8^; MC: t_139_ = 6.15, p = 10^-9^; CA3: t_103_ = 3.02, p = 0.003; CA1: t_144_ = 1.65, p = 0.1). During DS_L_, the discharge of GC-associated (n = 96) and MC-associated (n = 89) narrow-waveform interneurons reduces by 10% and 8%, respectively (Fig. 6C; GC In: t_95_ = 2.65, p = 0.009; MC In: t_88_ = 2.00, p = 0.05) while discharge of CA3-associated (n = 102) and CA1-associated (n = 148) narrow-waveform interneurons increases by 16% and 9%, respectively (Fig. 6C; CA3 In: t_101_ = 3.72, p = 0.0003; CA1 In: t_147_ = 2.40, p = 0.017). In contrast, during DS_M_, firing rates of GC-, MC-, CA3- and CA1-associated interneurons increase by 263%, 58%, 71% and 25% respectively (Fig. 6C; GC In: t_95_ = 9.51, p = 10^-15^; MC In: t_88_ = 3.41, p = 0.0009; CA3 In: t_101_ = 6.28, p = 10^-9^; CA1 In: t_147_ = 4.39, p = 10^-5^).

**Figure 6.**
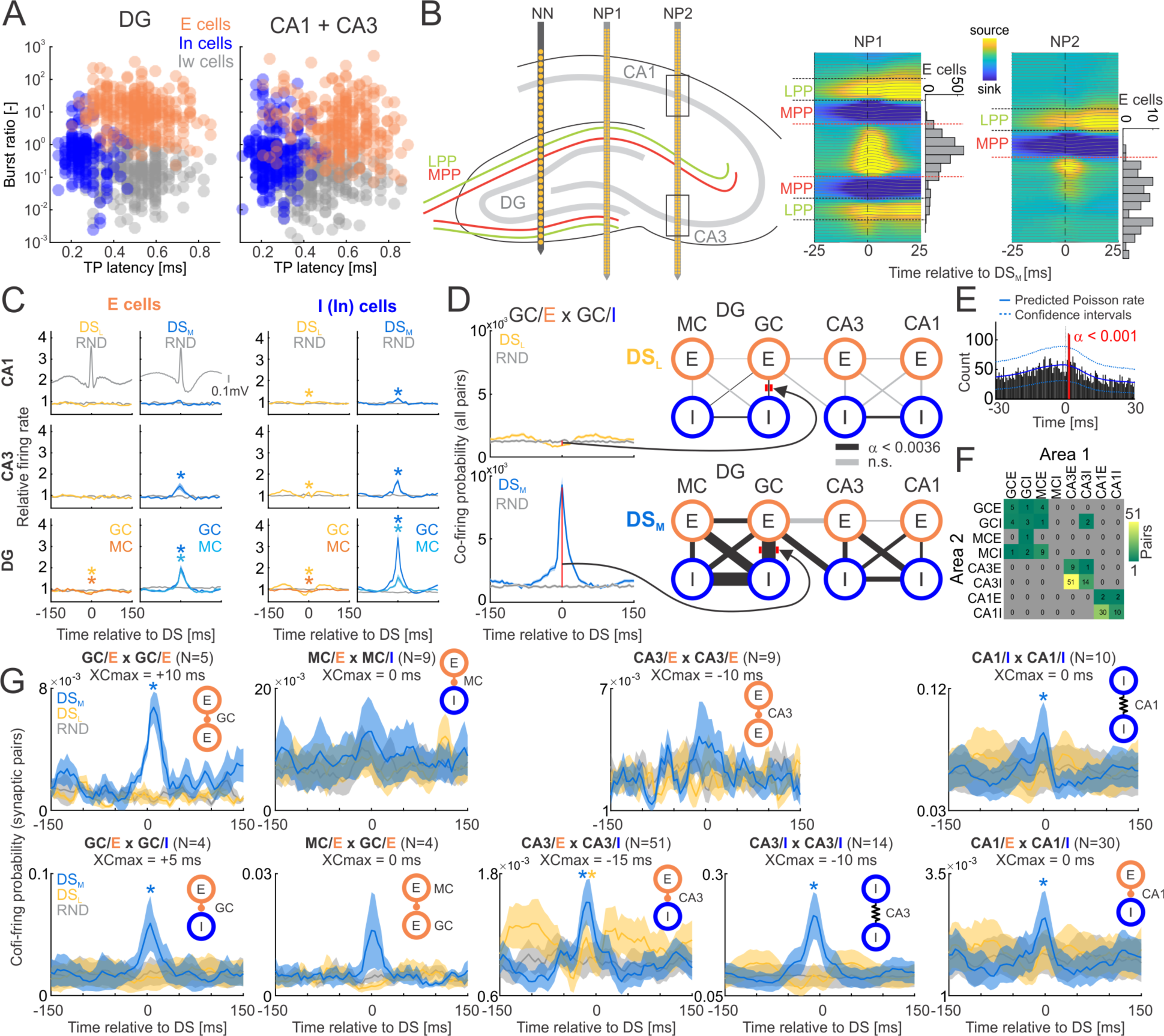
DS_M_ increases action potential discharge and cofiring in DG, CA3, and CA1 networks. A) Unitary action potentials were classified as being from excitatory cells (E), narrow-waveform interneurons (In) and wide-waveform interneurons (Iw) using the K-means algorithm on the DG-localized (left) and separately, the CA3- and CA1-localized datasets (right). B) Schematic mouse hippocampus (left) with medial placement of Neuronexus linear electrode array (NN) for detection of DS events and two example lateral placements of Neuropixels probes, NP1 proximal to DG, NP2 proximal to CA3. Distinctive DS_M_-triggered average CSDs distinguishes DG and CA3 localization. Rectangles along NP2 show CA1 and CA3 unit localization (Fig. S5A). Corresponding depth distributions of putative excitatory cells are shown on the right of CSDs. Dotted lines: DG borders of granule cell (red) and outer molecular (black) layers. C) Normalized firing rates of CA1 (top), CA3 (middle), and DG (bottom) units during DS_L_ (left) and DS_M_ (right) and random (gray) events. DS contaminated by SWR events were excluded. DS averages in gray. D) Representative cofiring probability of pairs of GC principal cells (GC/E) and proximal narrow waveform interneurons (GC/I) around DS_L_ (left, top, yellow) and DS_M_ (left, bottom, blue) and random (gray) events. Ratio of cofiring probability during DS events and random times is represented as line thickness (right) in a DG→CA3→CA1 network schematic; black connections significantly differ from random times, while grey connections do not after Bonferroni corrections (Table S1). E) Identification of a putative monosynaptic connection using enhanced spike-transmission strength in spike-time cross-correlogram. Solid blue line is expected cross-correlogram from Poisson model, dotted blue lines indicate confidence intervals, red bins mark significant deviations from the model. F) Summary matrix of counts of all identified monosynaptic pairs. G) Cofiring probability during DS_L_ (yellow), DS_M_ (blue) and random times (gray) in identified pairs with monosynaptic spike-transmission statistics. Title of each subplot shows time of maximum cross-correlation relative to DS, when statistics were computed. Stars mark significant deviations from random times. Averages ± S.E.M. are plotted.

These findings suggest that DS_L_ events result in net inactivation of both excitatory and inhibitory cells in DG and weak activation of interneurons in CA3 and CA1, without changing the firing rates of CA3 and CA1 principal cells, whereas DS_M_ events result in strong activation of both excitatory and inhibitory cells along the DG→CA3→CA1 trisynaptic pathway, with the primary effect in CA1 being to activate presumptive interneurons. The DS_M_-associated increase of both excitatory and inhibitory cells establishes conditions for enhanced temporal control of principal cell discharge through excitation-inhibition coordination, and enhanced opportunities for cofiring that can enhance neural transmission across the DG→CA3 →CA1 trisynaptic pathway (Ashhad and Feldman, 2020; Renart et al., 2010).

The hypothesis that DS_M_ promotes SG_dom_ by increasing neural control via CA3, predicts increased excitatory-inhibitory co-firing during DS_M_ (Ashhad and Feldman, 2020; Renart et al., 2010), as does a recent finding of increased cofiring between excitatory and inhibitory dentate cell pairs during moments of active and successful discriminative memory recall (van Dijk and Fenton, 2018). We analyze the cofiring of pairs of principal cells (E) and narrow-waveform interneurons (I) within ±3 ms of DS_L_, DS_M_ and random events (Fig. 6D; Table S1); DS events contaminated by SWR events were excluded. During DS_M_, cofiring amongst the GC and associated interneuron populations increases relative to chance (Fig. 6D, left; 649 ± 734%, t_464_ = 11.23, p = 10^-26^), whereas the cofiring decreases during DS_L_ (65 ± 108%, t_464_ = 4.89, p = 10^-6^). Principal cell and interneuron cofiring also increases during DS_M_ within CA3 (232 ± 299%, t_518_ = 7.18, p = 10^-12^) but does not change during DS_L_ (88 ± 87%; t_518_ = 2.26, p = 0.02). Increased cofiring between principal cells and interneurons is also observed within CA1 during DS_M_ (132 ± 137%; t_361_ = 5.46, p = 10^-7^) indicating potentially increased inhibitory control of principal cell spiking during DS_M_ but not during DS_L_ (105 ± 131%; t_361_ = 2.83, p = n.s. after Bonferroni correction). Cofiring also increased during DS_M_ but not DS_L_, between MC-and GC-associated interneurons (623 ± 676%, t_121_ = 8.90, p = 10^-15^) and between CA3- and CA1-associated interneurons (180 ± 214%, t_196_ = 5.53, p = 10^-8^).

These increases in the propensity for cofiring during DS_M_ imply that local neurotransmission between excitatory and inhibitory cells is enhanced between synaptically-coupled cell pairs during DS_M_. Enhanced spike-transmission strength estimated from cell pair spike time cross-correlograms has been used to identify monosynaptically (excitatory) coupled cell pairs (Fig. 6E; English et al., 2017; Stark and Abeles, 2009). Summary of the types of cell-class pairs identified by enhanced short-latency spike-transmission strength highlights a greater likelihood of detecting intra-regional coupling, including via common input, and electrical synapses (review Traub et al., 2018; Fig. 6F), as may be the case for interneuron-interneuron cell pairs that exhibit zero-lag coupling. The average cross-correlograms confirm that during DS_M_, cofiring is enhanced between excitatory – inhibitory cell pairs that are likely to be monosynaptically connected and possibly involved in rhythmogenesis (Figs. 3E, F; 4A) in DG, CA3 and CA1 (5 ms window; paired t test calculated at maximum cofiring value; GC/E x GC/I: DS_M_: t_3_ = 3.41, p = 0.04; DS_L_: t_3_ = 0.71, p = 0.53; CA3/E x CA3/I: DS_M_: t_50_ = 4.62, p = 10^-5^; DS_L_: t_50_ = 2.96, p = 0.004; CA1/E x CA1/I: DS_M_: t_29_ = 4.05, p = 0.0003; DS_L_: t_29_ = 1.43, p = 0.16). Furthermore, cofiring is enhanced during DS_M_ but not DS_L_ between pairs of granule cells (GC/E x GC/E; DS_M_: t_4_ = 5.99, p = 0.004; DS_L_: t_4_ = 1.11, p = 0.32), pairs of CA3 interneurons (CA3/I x CA3/I; DS_M_: t_13_ = 4.49, p = 0.0006; DS_L_: t_13_ = 0.37, p = 0.72) and pairs of CA1 interneurons (CA1/I x CA1/I; DS_M_: t_9_ = 3.68, p = 0.005; DS_L_: t_9_ = 1.61, p = 0.14). Taken together, DS_M_ selectively activates local excitation-inhibition network discharge in both DG and CA3, which control neuron cofiring between the DG and CA3 networks, likely to promote SG_dom_ and increased excitation-inhibition discharge in CA1.

Finally, because DS_M_ promotes synchronization of SG*_SR_* and SG_LPP_ in the slow gamma frequency range (Figs. 5C-F), neuronal discharge (Fig. 6C) and cofiring (Figs. 6D, G) within the DG-CA3-CA1 networks, it predicts that the DS_M_-enhanced SG*_SR_* gamma rhythm orchestrates the discharge through spike-field coupling that can maximize the efficiency of information transfer from DG to CA1. To evaluate this hypothesis, we examine the SG*_SR_* and SG_LPP_ spike-field coupling during DS events (Fig. 7). The spiking of dentate granule cells, CA3 and CA1 principal “E” cells is more organized at the trough of SG*_SR_* oscillations in CA1 during DS_M_ compared to DS_L_ (Figs. 7A-C; Kuiper test comparing the DS_L_- and DS_M_-associated discharge probability distributions across SG*_SR_* phase at time of the DS peak; GC/E: k = 1652, p = 0.001; CA3/E: k = 465, p = 0.02; CA1/E: k = 836, p = 0.01). In contrast, at the time of DS_M_, SG_LPP_ oscillations organize the local spiking of dentate GCs but not CA3 and CA1 principal cells when compared to DS_L_ (Figs. 7D-F; GC/E: k = 1508, p = 0.001; CA3/E: k = 374, p = 1; CA1/E: k = 516, p = 1). Similar relationships were observed for interneurons recorded in the vicinity of DG granule cells, CA3 and CA1 interneurons (Fig. S6). Taken together, these findings indicate that DS_M_ synchronizes discharge across the DG-CA3-CA1 trisynaptic circuit to SG*_SR_*, which enhances DG-CA1 transmission and promotes SG_dom_.

**Figure 7.**
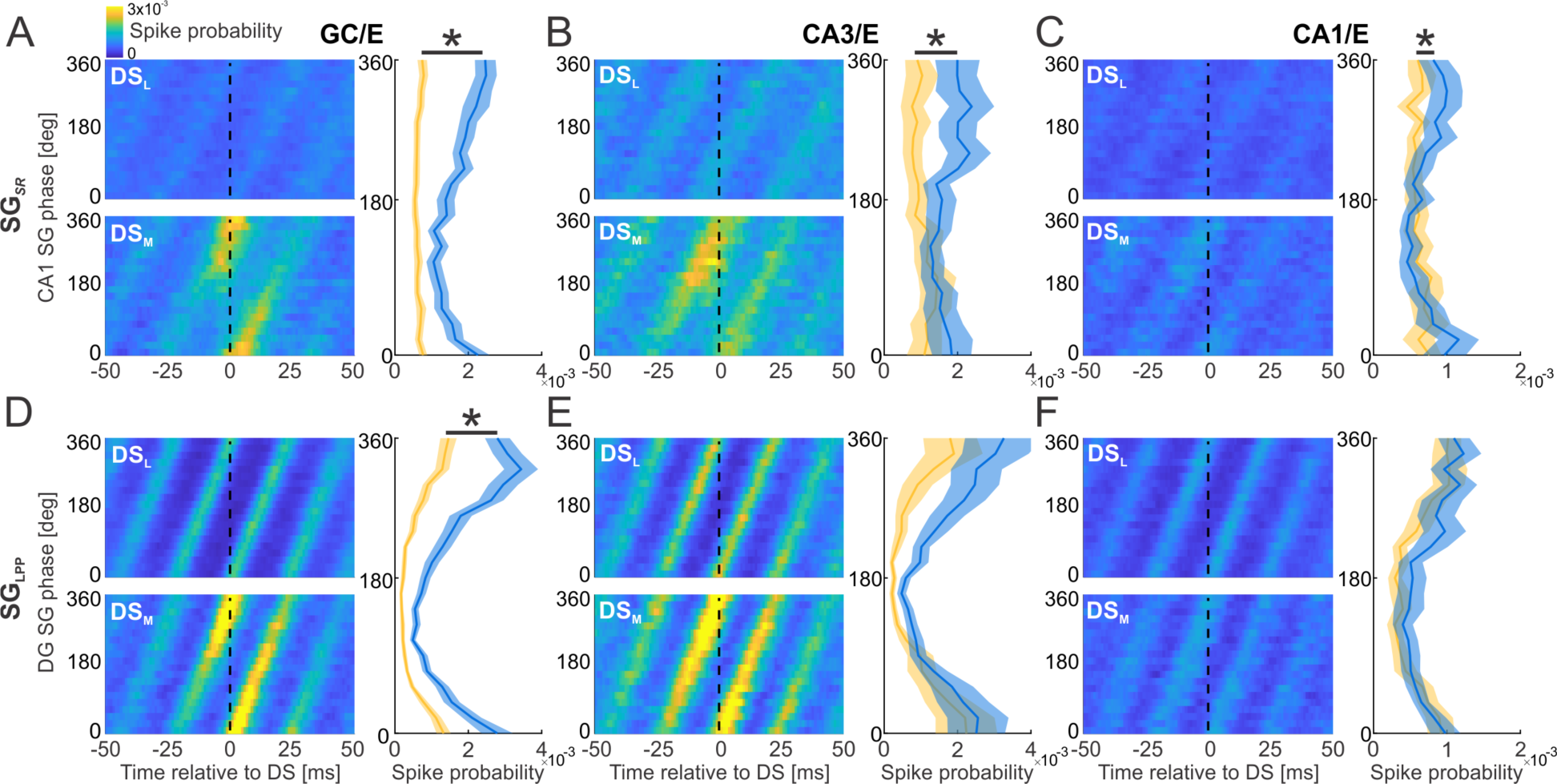
DS_M_ phase synchronizes discharge of GC, CA3 and CA1 cells through SG*_SR_*. A) Average discharge probability of a granule cell relative to DS times (x axis; DS_L_ top; DS_M_ bottom) and SG*_SR_* phase (y axis). Data averaged across cells for DS_L_ (yellow) and DS_M_ (blue) at DS peak (T = 0) shown on right. Stars mark a significant difference between the DS_L_ and DS_M_ phase distributions. Same as (A), but for B) CA3 and C) CA1 principal cells. D) Same as (A), but for SG_LPP_ gamma phase (y axis). Same as (D), but for E) CA3 and F) CA1 principal cells.

## Discussion

### DS_M_ control of information processing in Ammon’s horn

Our findings demonstrate entorhinal cortical control of information processing in hippocampus mediated by DS_M_, the result of the synchronous activation of medial perforant path terminals at the middle molecular layer of dentate gyrus (Fig. 2; Bragin et al., 1995). The effects of DS_M_ on the DG→CA3→CA1 network are in dramatic and consistent contrast to the effects of DS_L_ (Figs. 3-7), making it essential to distinguish them. Indeed, conclusions based on work that did not discriminate DS_L_ from DS_M_, have been hard to interpret (Bramham, 1998; Nokia et al., 2017).

We have even observed that place avoidance training causes synaptic plasticity of the MPP synaptic response in the suprapyramidal molecular layers of DG with a corresponding change of DS_M_ but not DS_L_, corroborating the two pathways are distinctive and can be altered independently by experience (Chung et al., preprint).

The present findings point to a process of dynamic control of hippocampal information processing marked by transient physiological events across the trisynaptic pathway. During SG_dom,_ DG, CA3, and CA1 discharge is transiently elevated along with DG-CA3 cofiring (Fig. 6), and there is slow gamma frequency and phase synchronization between LPP inputs to DG and the *stratum radiatum* input from CA3 to CA1 (Figs. 5, 7), indicating involvement of the entire trisynaptic pathway, similar to SWR (Buzsaki et al., 2003; Sullivan et al., 2011). Indeed, during DS_M_, SWR probability in a 20 ms window increases from 1.2% to 1.8% (Fig. 3G) and place cell discharge is non-local during SWR (Buzsaki, 2015; O’Neill et al., 2006; Papale et al., 2016; Sullivan et al., 2011) as well as during SG_dom_ (Fig. 1; Dvorak et al., 2018).

The qualitative distinction between information signaled by LEC and MEC is important in this context. LEC transmits contextual information based on objects and egocentric information that constitutes the content of spatial experience (Knierim et al., 2014; Tsao et al., 2013; Wang et al., 2018), whereas MEC transmits allocentric spatial signals like direction, distance, borders, and speed (Hargreaves et al., 2005; Rowland et al., 2018; Sargolini et al., 2006; Ye et al., 2018). Remarkably, the MEC-originating DS_M_ signal that promotes SG_dom_ and switches CA1 to non-local positional memory processing is coincident with synchronization between CA3-transmitted slow gamma inputs to CA1 and LEC-transmitted slow gamma inputs to DG, rather than from MEC (Figs. 5, 7). Within the “communication through coherence” hypothesis (Fries et al., 2007), the LEC→DG and CA3→CA1 inputs have a privileged opportunity for information transfer via synchronized slow gamma transmission, and so we speculate that perhaps while switching the hippocampal information processing mode, DS_M_ loads Ammon’s horn with information from the LEC-delivered egocentric contents of experience. In this way, during DS_M_, the consequent activation of CA3 may be preferentially influenced by contextual and egocentric, ecphoric retrieval cues delivered by the LEC inputs (Kelemen and Fenton, 2013; Tulving et al., 1983). If these manifest as SG_dom_ because of the strong DS_M_-associated activation of CA3 (Figs. 6, 7), the result is control of CA1 output that overrides the default control that is exerted by the mid-frequency gamma mediated ECIII input at *slm* (Brun et al., 2002). These *slm* inputs appear necessary for place cell firing (Brun et al., 2008), and they create permissive dendritic depolarization (Jarsky et al., 2005), but in principle can be shunted by the slow gamma-associated inputs (Keeley et al., 2017), and are possibly actively inhibited as a consequence of DS_M_ (Fig. 4F), all of which promote SG_dom_ (Fig. 3H). As we have observed during SG_dom_, CA1 discharge will signal non-local positions that during a memory task, correspond to recollected places (Dvorak et al., 2018), and we observed here a correspondingly reduced local discharge by place cells in their firing fields (Fig. 1), despite maintained CA1 firing (Fig. 6C). Indeed, the findings of a hippocampus-wide (Figs. 6, 7), DS_M_-promoted SG_dom_ change to a non-local mode of information processing identifies a source of the overdispersion that is characteristic of place cells in CA1, CA3, and DG (Fenton et al., 2010; Fenton and Muller, 1998; Hok et al., 2012; Jackson and Redish, 2007; van Dijk and Fenton, 2018), and also grid cells, although we cannot conclude the mechanism is the same (Nagele et al., 2020). The findings also offer an explanation for the possible utility of CA1 receiving two spatial inputs; the Schaffer collaterals provide place cell inputs that can be non-local and related to mental experience, whereas the temporoammonic pathway provides an input comprised of components of place (grid cell distances, directional cells, border cells, and speed cells) more tethered to local, physical experience.

## STAR Methods

### Subjects

A total of 14 wild-type mice with a mixed C57BL/6J background were used for the study. The mice were 3 - 6 months old during surgery. Nine mice were implanted with linear silicon arrays. Three mice were implanted with a metal head plate for head-fixed recording using linear silicon arrays and Neuropixels probes (Jun et al., 2017). Two mice from a previously published dataset were implanted with tetrodes (Dvorak et al., 2018).

### Surgery

LFP recordings were collected using 32-channel (8 mice) and 16-channel (1 mouse) linear silicon array electrodes (Neuronexus, Ann Arbor, MI) with 50 µm spacing and 703 um_2_ electrode area. The 32-channel electrodes spanned both CA1 and DG locations, and the 16-channel electrode spanned only DG locations. The electrodes were implanted stereotaxically under isoflurane anesthesia (2%, 1 L/min). The tip was aimed at -1.85 AP, ±1.20 ML, -2.3 DV relative to bregma. The electrodes spanned the dorso-ventral axis of the dorsal hippocampus.

Reference electrodes were aimed at the cerebellar white matter. The electrode assemblies were anchored to the skull using 3-4 bone screws with dental cement (Grip Cement, Dentsply, Milford DE). One anterior screw was used as a ground. A four-wire stimulating electrode bundle was made by twisting together four 75-µm diameter nichrome wires (California Fine Wire, Grover Beach, CA). The bundle was cut at an angle so as to span 0.5 mm. During surgery, the stimulating bundle was placed in the ipsilateral perforant path +0.5 AP, ±4.1 ML, 1.0-1.6 DV from lambda. Evoked response waveforms were carefully checked with different pair combinations of stimulation electrode channels. In the mice that were used for head-fixed recordings, a titanium head plate was attached to the skull using dental cement and the exposed skull was covered with KwikSil, a low toxicity adhesive (World Precision Instruments,

Sarasota, FL) and protected by attaching a plastic cup. All mice were allowed at least 1 week to recover. In mice that were used for head-fixed recordings, a secondary surgery was performed immediately before the experiment. The plastic cup and KwikSil were removed and a craniotomy was made at 1.85 AP, ±1.20 ML relative to bregma to enable electrode placement. Between consecutive days of recordings, a KwikSil protective cup assembly was reattached to prevent infection.

### Behavioral tasks

Nine mice with implanted linear silicon probe arrays and 2 mice implanted with tetrodes were trained in the active place avoidance task. Each day consisted of a 30 min rest session in the home-cage, which was placed in the recording chamber, followed by a 30-min place avoidance session. After habituation (shock off), a total of 3 training sessions (shock on) were administered to all animals, one session each day. A retention session with the shock on followed 1 week after. Electrophysiology data from 9 mice implanted with linear silicon probe arrays and recorded during rest were used for Figs. 2, 3A-G and 5. Electrophysiology data from the same group recorded during place avoidance were used for Figs. 1A, B, 3H and 4. Electrophysiology data from the 2 mice implanted with tetrodes were used for Figs. 1C, D. Two mice were recorded in a custom head-fixed setup using both Neuropixels and Neuronexus linear silicon probe arrays during 59 sessions (average length 15.7 ± 3.9 min) spread over multiple days.

Mice were encouraged to walk using a custom water delivery system. Electrophysiology data recorded in head-fixed mice were used for Figs. 6, 7.

### Electrophysiology recording

An amplifier board with 32 unipolar inputs and 3-axis accelerometer (RHD2132, Intan Technologies, Los Angeles, CA) was connected directly to the Neuronexus probe for signal amplification and digitization. A lightweight, counter-balanced cable (Intan Technologies, Los

Angeles, CA) was used to power the amplifier board and the infrared LED used for tracking as well as to transmit digital data to the computer using a custom recording system connected to the USB port of a PC. The cable was connected through a lightweight commutator to enable free movement of the animal. The signal from each electrode was low-pass filtered (500 Hz) and digitized at 2 kHz. Evoked responses were obtained using a constant current stimulus isolation unit (WPI, Sarasota, FL; model: A365RC) that was used to deliver individual unipolar 100 µs stimulus pulses across the electrode pair. Evoked responses were low-pass filtered at 4 kHz and sampled at 8.12 kHz. In head-fixed recordings, the signal from a Neuropixels probe was filtered between 0.5 Hz and 1 kHz and sampled at 2.5 kHz for LFP recordings and filtered between 300 Hz and 10 kHz and sampled at 30 kHz for single unit recording. Both electrophysiology systems were synchronized using square TTL pulses generated by the Neuropixels system that was recorded by both systems. The animals were recorded during a 30-minute session in their home-cage during the first exposure to the experimental room. The mouse’s movements during recordings were monitored continuously using a video tracking system (Tracker, Bio-Signal Group, Acton, MA) that was synchronized to the electrophysiology data using the video frame pulses generated by the camera.

### Data Analysis

#### LFP and electrode localization

LFPs were localized by visual LFP inspection of sharp-wave ripples in CA1 *stratum pyramidale* and dentate spikes in the hilus of the dentate gyrus (Fig. S1E). Electrode locations were verified histologically at the end of recordings (Fig. S1F, G). In the mouse implanted with the 16-ch electrode array, only dentate LFPs were recorded because of the limited spatial span of the electrode.

#### Detection of SG_dom_ events

A published algorithm was used to extract oscillatory events from LFP or independent components obtained using ICA (Dvorak and Fenton, 2014). The LFP is transformed into a time-frequency power representation by convolving the LFP/IC signal with a group of complex Morlet wavelets and z-score normalizing each band-specific signal. Oscillatory events are detected as local peaks in the normalized 2-D time-frequency space. Detection of oscillation rates and SG_dom_ events was described previously (Dvorak et al., 2018). Oscillation rates (Fig. 1A, lower) are computed as the number of detected events in a representative frequency range (30-50 for CA1 slow gamma, 70-90 Hz for CA1 mid-frequency gamma) in a 1-s window advanced by 0.25 s and smoothed using 2.5 s moving average. SG/MG ratio (Fig. 1A, lower) is computed as a ratio of CA1 slow gamma oscillation rate and CA1 mid-frequency gamma oscillation rate. SG_dom_ events, are defined as local peaks in SG/MG ratio with prominence exceeding 1 and SG/MG ratio > 1 (corresponding to SG > MG).

#### Detection and classification of dentate spikes

The LFP channel with the largest visually identified amplitude of dentate spike was band-pass filtered 5-100 Hz and the amplitude was z-score normalized. Next, all local peaks in the band-pass signal were detected, and several features were extracted, including the amplitude difference between the DS maxima and the preceding as well as following minima, and also the spike width that was measured at the level of either the preceding or following minima, whichever was closer to the maxima. The spike amplitude distributions were further normalized by z-score normalization of the log-transformed amplitudes. Putative dentate spikes were selected if their prominence (difference between amplitude of the DS maxima and the smaller of either its preceding or the following minima) > 0.75 and when the width of the event was between 5 and 25 ms. The optimal values were selected based on analysis of the feature histograms.

Dentate spikes were classified as DS_L_ and DS_M_ based on their CSD profiles (Bragin et al., 1995). CSDs were calculated using the CSDplotter Matlab toolbox (Pettersen et al., 2006) at the peak of a putative DS event. Independently for each recording, the CSD for each putative DS event was analyzed for local minima, corresponding to CSD sinks (Fig. 2B). The histogram of detected local minima of all putative DS events was plotted and the local maxima that represented the highest probability of current sinks were manually identified (color dots in Fig. 2B, middle). This analysis resulted in 4 locations corresponding to (from top to bottom) the outer and the middle molecular layers of the suprapyramidal DG blade, and the middle and the outer molecular layers of the infrapyramidal DG blade. DS_L_ and DS_M_ were then identified from the suprapyramidal DG blade as putative DS events with a sink occurring ± 25 µm around the location selected in the sink histogram (solid lines in Fig. 2B, right), although average CSD profiles of classified DS events did not change when classification from infrapyramidal DG blade was used instead (dashed lines in Fig. 2B, right). Putative DS events with sinks in both the outer and middle molecular layers (1.7 ± 1.5 %; mean ± S.D.) as well as those with no sinks detected in either the outer or the middle molecular layers (30.5 ± 17.8 %) were excluded from analyses. Only putative DS events with a sink exclusively in either the outer or the middle molecular layer (67.8 ± 18.3 %) were classified as DS_L_ and DS_M_ respectively.

#### Detection of ripple events

We followed a previously published algorithm (Csicsvari et al., 1999) with several modifications to detect ripple events. We used the LFP recorded from the CA1 *stratum pyramidale* electrode, where ripples were identified visually. First, the signal was 150-300 Hz band-pass filtered. Next, we computed the sliding root-mean square (RMS) estimate in a 10-ms window. Next, we z-scored normalized the RMS estimate and detected the local maxima with z > 3. Finally, for each detected event, we computed the wavelet time-frequency representation of the LFP and for each detected event we extracted its frequency as a local peak in the time-frequency wavelet spectrum (similar to detection of gamma oscillations described earlier). Only events with frequencies between 130-250 Hz were selected for further analysis.

#### Independent components analysis of the LFP

We used independent component analysis (ICA) to extract the specific CA1 dendritic components (Fernandez-Ruiz and Herreras, 2013; Fernandez-Ruiz et al., 2017), which minimizes the impact of volume conducted signals and estimates the components that can be precisely matched to specific dendritic compartments. LFP signals that were recorded using linear silicon array electrodes were decomposed into individual dendritic components using a previously described procedure (Fernandez-Ruiz and Herreras, 2013; Fernandez-Ruiz et al., 2017; Makarov et al., 2010) with several modifications. First, LFP signals were filtered between 20 Hz and 150 Hz. Next, principal component analysis (PCA) was applied to the filtered LFP data in order to find out how many principal components explain over 99% of the signal variance in the data. Next, independent component analysis (ICA) was applied to the filtered LFP data using the FastICA Matlab toolbox (Hyvarinen, 1999) by specifying the number of principal components that were obtained in the previous steps for both PCA-based dimensionality reduction and the target number of resulting independent components. Next, components of the unmixing matrix were used to compute CSDs of the individual voltage loadings for component localization and independent components (ICs) were processed using comodulogram analysis for frequency-based classification of components. Here, we took advantage of theta phase coupling of gamma oscillations, which can reveal a specific frequency footprint of each component (Schomburg et al., 2014). Specifically, the LFP from the *stratum pyramidale* electrode was filtered using a set of FIR filters with 2 Hz bandwidth, in the range 5-11 Hz followed by the Hilbert transform to obtain the phase of CA1 theta oscillations. Next, independent components were filtered using 20-Hz wide filters in the range 20-150 Hz followed by the Hilbert transform to obtain amplitude information from individual components. Details of the filters and filtering procedure were described previously (Dvorak and Fenton, 2014). The phase and amplitude information were then combined between all pairs of frequency bands used to obtain phase and amplitude information and a modulation index (Tort et al., 2010) was computed for each pair resulting in a comodulogram (Figs. 3C, 5A) that reveals the peak coupling between the phase of theta and the amplitude of a given IC. We found that the ICA analysis provides better segregation of the independent components if the number of LFP channels is restricted before performing ICA. On the other hand, it is not possible to say which LFP channels to include in the analysis for best IC separation. Consequently, we performed a grid search, where we systematically repeated ICA for different numbers of included contiguous segments of LFP channels referenced either to *stratum pyramidale* for CA1 (Fig. 3) or to the *hilus* for DG (Fig. 5). The resulting CSD profiles of ICs were then visually compared and selected based on both the CSD profile of voltage loadings and a clearly isolated peak of coupling between theta phase and the amplitude of a given component. While this operation is extremely computationally intensive, it allowed robust detection of the corresponding components in all the mice we studied (Fig. S2A).

#### Phase locking analysis

To study the phase coupling between different oscillatory rhythms, we used the phase locking value (PLV) estimate (Lachaux et al., 1999), which provides a good estimate of phase locking for signals where the volume conducted signals have been minimized by ICA (Vinck et al., 2011). To calculate PLV of a pair of signals, we used an array of complex Morlet wavelets spaced by 1 Hz between 20 Hz and 100 Hz convolved with each of the ICs in the pair to obtain the instantaneous phase of both ICs at a given frequency. Next, we computed the instantaneous phase difference between the two ICs, IC_1_ and IC_2_. Then, for all pairs of time offsets in the range -100 ms to +100 ms relative to the DS event and each frequency, we computed instantaneous phase differences across all DS events Δφ(*t, f*) = φ_1_(*t, f*) − φ_2_(*t, f*). Finally, we computed PLV across DS events as 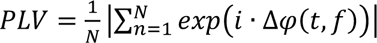, where *i* is the imaginary unit, N is number of DS events, t is the offset relative to DS event and f is frequency used to filter the signal to obtain its phase. Repeating this algorithm for a range of frequencies and offsets relative to DS events generates a time-frequency PLV estimate that is centered at each type of DS (Fig. 5C).

#### Detection of oscillatory cycles

To detect oscillatory cycles of oscillatory bursts (Fig. 4), we started by correcting the polarity of ICs, because the polarity of an individual ICs is arbitrary (Hyvarinen, 1999). Here, we took advantage of the known relationship between hippocampal gamma oscillations < 100 Hz and the spectral leakage of spiking activity (Fig. S2B), that can be observed > 150 Hz at *stratum pyramidale* (Dvorak and Fenton, 2014; Lasztoczi and Klausberger, 2016; Schomburg et al., 2012). We first created a set of Morlet wavelets covering 20-50 Hz for SG*_SR_* or covering 60-90 Hz for MG*_SLM_* and used them to obtain the instantaneous phase of the IC components at specific frequencies. Next, we filtered the LFP from *stratum pyramidale* in the frequency range 150-250 Hz followed by a Hilbert transform to obtain the amplitude of the high frequency activity that served as a proxy for spiking activity. Next, for each IC, we created a phase-amplitude histogram of 150-250 Hz amplitude distribution relative to the phase of the IC component either in the 20-50 Hz range or the 60-90 Hz range (Fig. S2B). Finally, we visually compared the resulting relationships and corrected each component so that the 150-250 Hz spiking-proxy activity was maximal at the descending phase of the SG*_SR_* close to the trough and the ascending phase of MG*_SLM_* close to the trough (Fig. S2B). This step allowed us to reliably correct the polarity of all components from all animals in order to reliably extract local minima of oscillatory bursts. After correcting the polarity of IC components, we detected oscillatory bursts as described earlier and then found local minima in the ± 50 ms window around an oscillatory peak for SG*_SR_* and in the ± 29 ms window around the oscillatory peak for MG*_SLM_* corresponding to 3 cycles of oscillatory activity centered at the oscillatory cycle with largest amplitude (Fig. S2C). The timestamps of individual oscillatory cycles were recorded and used for later processing.

#### Single unit analysis

Single units were sorted using a published open-source algorithm Kilosort2 (Pachitariu et al., 2016) that is optimized for Neuropixels probes and takes advantage of GPU processing to improve algorithm performance. After automated clustering of the data, we selected only units with < 20% estimated contamination rate with spikes from other neurons that were computed from the refractory period violations relative to expected. We also excluded units with non-characteristic or noisy waveforms resulting in identifying a total of 9404 single units.

The units were then localized to neocortex, CA1, DG/CA3 and thalamus using three criteria: 1) the depth of the Neuropixels probe relative to the cortical surface, 2) localization of dentate spikes in the hilus of dentate gyrus and sharp wave ripples in CA1 *stratum pyramidale* and clustering of units along the depth of the linear Neuropixels array. The clustering of units into different regions becomes apparent when we plot the depth of the maximal amplitude of the average action potential waveform for each unit along the length of the probe (Fig. S5A). The cluster of single units that overlaps with the detected location of sharp-wave ripples was classified as CA1, whereas the cluster of units that overlaps with the detected location of DS was classified as DG/CA3. The cluster of units between CA1 and the cortical surface was classified as neocortical neurons and the large amplitude units below DG/CA3 were classified as thalamic neurons. To separate CA3 from DG units, we used two additional criteria. First, we used the anatomical location, confirmed by histology, and considered CA3 units to only be from electrodes that were more lateral than +1.5 mm relative to the midline. Second, we took advantage of the asymmetric profile of the perforant path termination in DG that is apparent in the CSD profiles of LFPs that were recorded with Neuropixels probes and triggered by DS_M_ events (Fig. 6B). ECII projections to the infrapyramidal molecular layers of DG terminate at the mediolateral extent at which CA3 begins, while ECII projections to the suprapyramidal molecular layers of DG continue in parallel with CA3 (Fig. 6B; Fig. S1F). Electrodes that exhibited only a dorsal current sink were classified as CA3, while electrodes that exhibited a symmetrical pair of current sinks were classified as DG. To further classify DG cells as putative granule cells (GC) and mossy cells (MC) we took advantage of two identified locations, that of the granule cell layer at the CSD reversal between the current sink in the middle molecular layer and the current source in the hilus triggered by DS_M_, and that of the maximal amplitude of the average action potential of a given cell. Cells within 150 μm of the CSD reversal were classified as GC, while cells deeper than 150 μm were classified as MC (Senzai and Buzsaki, 2017). This procedure resulted in localizing 1413 cells to neocortex, 6422 neurons to thalamus, 492 cells localized to CA1, 696 cells localized to DG and 285 cells localized to CA3.

To classify units into putative excitatory and inhibitory neurons we used a similar approach as in other studies (Jia et al., 2019; Senzai and Buzsaki, 2017; Talbot et al., 2018) and extracted several features associated with the average action potential waveshape and features associated with firing properties (Fig. S5B). Datasets were split into DG cells and CA1 + CA3 cells because features of DG action potentials were visually different from those in CA1 + CA3 (Fig. S5C). Consequently, the two datasets were independently analyzed using the k-Means algorithm implemented in JMP 14 software to identify three clusters corresponding to three types of neurons classified as principal cells (E), narrow-waveform interneurons (In) and wide-waveform interneurons (Iw). Classification of CA1+CA3 cells separately from DG cells led to the best classification results into the selected neuronal subtypes. In the analyses that follow, we only focus on E and In cells because of their maximal separation in the feature space (Fig. 6A).

#### Peri-DS-event time cofiring histogram

We assessed the probability that a pair of cells would cofire relative to the occurrence of a dentate spike by computing a cofiring probability for each cell pair. The probability was computed in a 6 ms-long window centered on the dentate spike peak. The co-firing probability was compared to randomly sampled events to obtain a ratio of cofiring change. Statistical validation was computed using a t test between the cofiring probabilities during DS events and randomly sampled times. The significance threshold was corrected using Bonferroni’s method.

#### Bayesian location decoding

To obtain estimates of the mouse’s location based on single unit data, we used a published algorithm (Zhang et al., 1998), where the probability of the current location is defined as 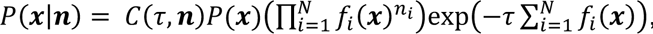 where *C*(*τ,n*) is a normalization factor so that ∑_*x*_*P*(*x*|*n*) = 1, *f*_*i*_(*x*) are firing rate maps for cells *i..N* obtained either by binning the 2-D space into 32x32 bins (Fig. 1C) or 1-D space (distance to shock zone) into 12 angular bins (Fig. 1D), *P*(*x*) is the dwell distribution, *τ* is the length of the time window (500 ms), *n*_i_ is the number of spikes fired by the i-th cell in a given time window and ***x*** is the *(x,y)* position of the animal in the 2D analysis or the angular position in the 1D analysis. Only recordings with at least five high quality spatial or non-spatial putative pyramidal cells were analyzed. Time windows with no spikes were excluded from analysis. Decoded location probability during SG_dom_ (Fig. 1D) was normalized by a decoded location probability during MG_dom_ (SG_dom_ functional counterpart), computed as local peaks in the ratio of CA1 mid-frequency gamma and CA1 slow gamma). Statistical analyses were performed using JMP version 14 (SAS, Cary, NC) and Matlab 2019b (Mathworks, Natick, MA). Significance was accepted at p < 0.05. Exact p values are reported throughout.

