## Supplemental Information for "Dentate spikes and external control of hippocampal function"

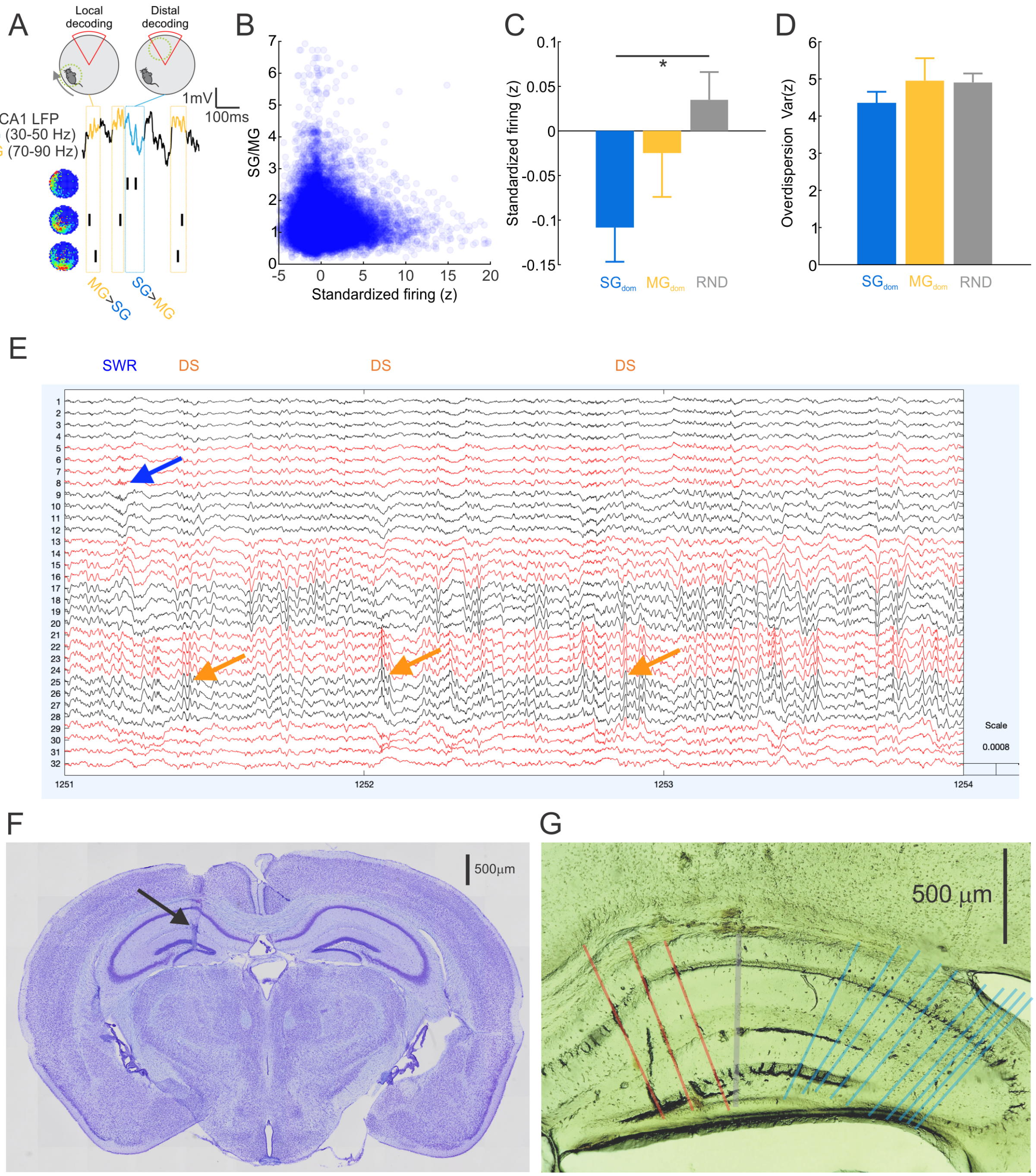


**Supplemental figure S1 related to Figures 1 and 2.** A) Schematic with raw LFP and spiking from three place cells illustrate that during slow gamma-dominant episodes (SG/MG>1; blue), decoding of CA1 place cells is non-local, and points to remote locations associated with the shock zone. During mid-frequency gamma-dominant episodes (MG/SG>1; yellow), the decoding is local, and points to the mouse's current location. B) Relationship between SG/MG ratio and standardized firing ($z$) demonstrating that place cells discharge less than expected when mice walk across a cell's place field during $SG_{dom}$. This is because the firing is non-local. Notice lower z values during larger SG/MG corresponding to $SG_{dom}$. Recordings were split into 5-s long episodes and for each episode we calculated the expected number of spikes as $EXP = \sum_i r_i * t_i$, where $r_i$ is the time-averaged rate at location i, and $t_i$ is the time spent in location i during the episode. For each 5-s interval, we then calculated z, the standard normal deviate of the observed discharge ($OBS$) computed under the inhomogeneous Poisson assumption as $z = \frac{OBS-EXP}{\sqrt{EXP}}$. C) Standardized firing ($z$) during $SG_{dom}$ (peaks in SG/MG ratio with prominence > 1), $MG_{dom}$ (peaks in MG/SG ratio with prominence > 1) and random events. Standardized firing (z) was lower during $SG_{dom}$ events than the other events (ANOVA: $F_{2,521} = 3.16$, $p = 0.04$, $SG_{dom} > MG_{dom}$ = RND). D) Overdispersion - variance of standardized firing [Var(z)] measuring the reliability of spatial firing during $SG_{dom}$, $MG_{dom}$ and random events when *EXP* > the cell's average rate, as this selects passes through the cell's firing field. The variance of the distribution of the standardized firing, z describes overdispersion. Overdispersion was similar across the three conditions and comparable to previously published values (ANOVA: $SG_{dom}$ = 4.25; $MG_{dom}$ = 4.80; RND = 5.11; $F_{2,521} = 1.03$, $p = 0.36$). E) Localization of DS events (*hilus*: channel 25; orange arrows) and a SWR event (*stratum pyramidale*: channel 8; blue arrow) in 3 seconds of LFP data. Scale (Volts) shown on right. F) Photomicrograph showing the trace (black arrow) of a Neuronexus 1x32 silicon electrode array (6 mm shank length, 50 µm inter-site spacing, 703 µm$^2$ site area. G) Photomicrograph showing traces of a Neuronexus probe (gray) and Neuropixel probes targeting DG (red) and CA3 (blue) using CM-DiI dye (ThermoFischer catalog V7001).

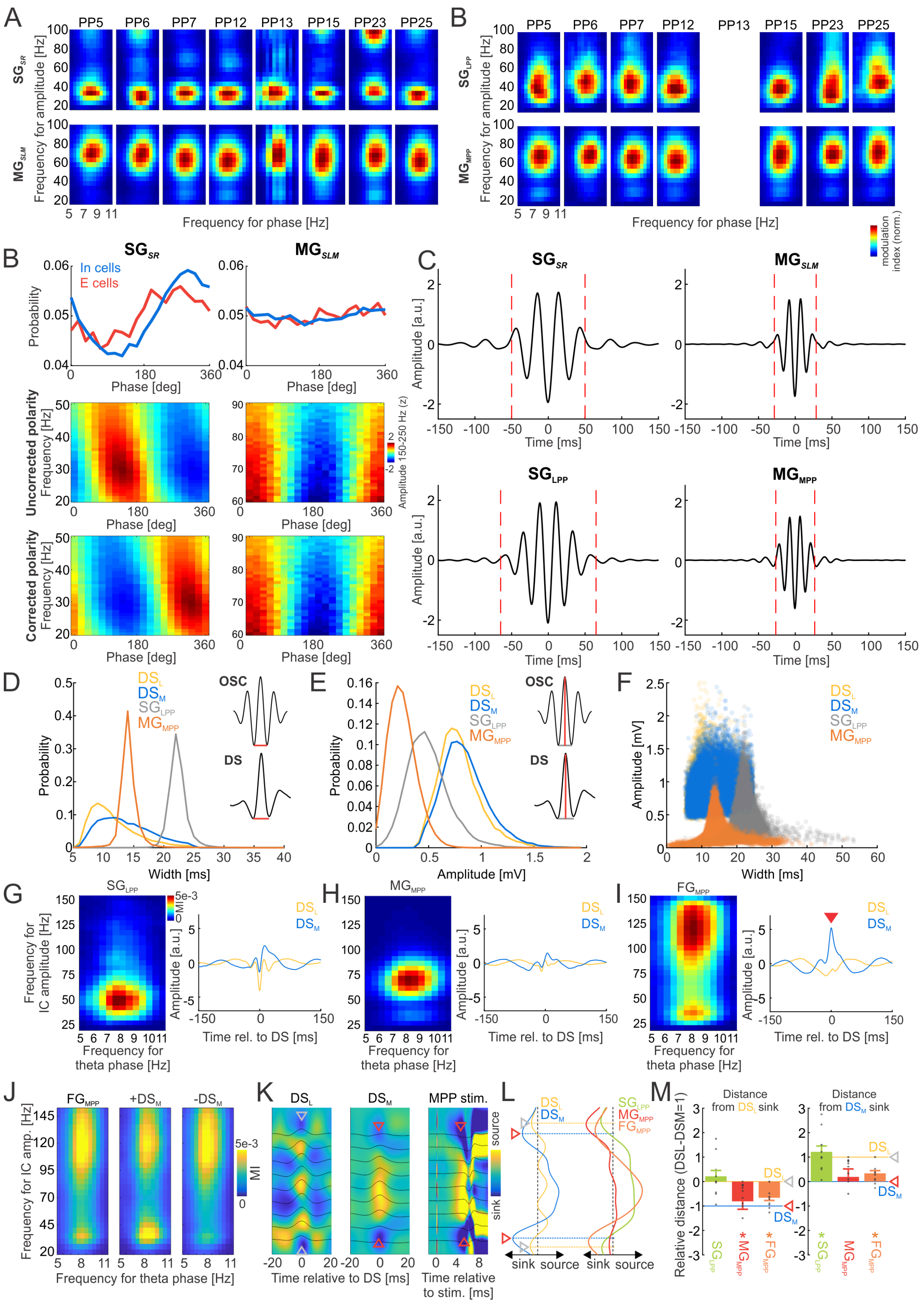
A
PP5 PP6 PP7 PP12 PP13 PP15 PP23 PP25
SG_SR
MG_SLM
Frequency for amplitude [Hz]
Frequency for phase [Hz]
B
SG_LPP
MG_MPP
modulation index (norm.)
In cells
E cells
Probability
Phase [deg]
Uncorrected polarity
Corrected polarity
Frequency [Hz]
Amplitude 150-250 Hz (z)
C
Amplitude [a.u.]
Time [ms]
D
DS_L
DS_M
OSC
DS
Width [ms]
E
Amplitude [mV]
F
G
H
I
FG_MPP
Frequency for IC amplitude [Hz]
Frequency for theta phase [Hz]
Time rel. to DS [ms]
J
+DS_M
-DS_M
Frequency for IC amp. [Hz]
MI
K
MPP stim.
Time relative to DS [ms]
Time relative to stim. [ms]
source
sink
L
sink source sink source
M
Distance from DS_L sink
Distance from DS_M sink
Relative distance (DSL-DSM=1)

**Supplemental figure S2 related to Figures 3 and 4.** A) Comodulograms between *stratum pyramidale* theta (5-11 Hz) LFP phase (horizontal axis) and amplitude of isolated independent components (20-100 Hz; vertical axis) for CA1 slow gamma ($SG_{SR}$; *stratum radiatum* IC; top) and CA1 mid-frequency gamma ($MG_{SLM}$; *stratum lacunosum moleculare* IC; bottom). B) Comodulograms between *stratum pyramidale* theta (5-11 Hz) LFP phase (horizontal axis) and amplitude of isolated in dependent components (20-100 Hz; vertical axis) for DG slow gamma ($SG_{LPP}$; lateral perforant pathway IC; top) and DG mid-frequency gamma (medial perforant pathway IC; bottom). Each panel corresponds to a single animal. B) Probability of discharge of putative CA1 excitatory cells (E, red) and narrow waveform interneurons (In, blue) in relation to the phase of the CA1 slow gamma ($SG_{SR}$; top left) CA1 mid-frequency gamma ($MG_{SLM}$; top right). Uncorrected CA1 slow gamma (middle left) and mid-frequency gamma (middle right) phase modulation of *stratum pyramidale* high frequency (150-250 Hz) LFP amplitude selected as a proxy of spiking. Notice reversed polarity in case of $SG_{SR}$ but correct polarity in case of $MG_{SLM}$. Corrected polarity using manual component polarity inversion (bottom). Notice the alignment of discharge preference (top) and amplitude of the high frequency *stratum pyramidale* LFP (bottom). C) Average oscillatory event of CA1 slow gamma ($SG_{SR}$; top, left), CA1 mid-frequency gamma ($MG_{SLM}$; top, right), DG slow gamma ($SG_{LPP}$; bottom, left) and DG mid-frequency gamma ($MG_{MPP}$; bottom, right). All oscillatory events were time-locked to local minima localized closest to the maximal oscillatory power. Oscillatory events were averaged across all animals. Red lines indicate borders of oscillatory events resulting in duration of 100 ms for $SG_{SR}$, 57 ms for $MG_{SLM}$, 130 ms for $SG_{LPP}$ and 53 ms for $MG_{MPP}$. Only oscillatory cycles detected within the identified borders were used in analyses. D) Probability distributions of widths of positive hilar LFP waveforms identified at times of detected $SG_{LPP}$ and $MG_{MPP}$ oscillatory events (OSC; inset) and measured between local minima preceding and following the largest amplitude of DS events (DS; inset). Notice different probability distributions of $DS_L$/$DS_M$ and $SG_{LPP}$ and $MG_{MPP}$ oscillatory events, demonstrating robustness of separation of the two classes of events, even though the LPP projection from LECII is the origin for both $DS_L$ and $SG_{LPP}$ and the MPP projection from MECII is an origin for both $DS_M$ and $MG_{MPP}$. E) Probability distributions of amplitudes of positive hilar LFP waveforms identified at times of detected oscillatory events (OSC; inset) and measured as average voltage between the largest amplitude and the preceding and following local minima of DS events (DS; inset). Notice different probability distributions of DS and oscillatory events, again demonstrating robustness of separation of the two classes of events. F) Scatterplot of width vs. amplitude of detected DS events and amplitudes of detected positive LFP waveforms detected during oscillatory events. Width and amplitude features of positive hilar LFP waveforms associated with $SG_{LPP}$ events and $DS_L$ overlap in 4.77% of events, $SG_{LPP}$ and $DS_M$ overlap in 10.38% of events, $MG_{MPP}$ and $DS_L$ overlap in 15.16% of events and $MG_{MPP}$ and $DS_M$ overlap in 17.36% of events. Those overlaps are further reduced by CSD-based $DS_L$ and $DS_M$ classification, where only DS events with precise location of CSD sinks in the outer and middle DG molecular layers, respectively, are selected (Fig. 2B). G) Comodulogram between the phase of *stratum pyramidale* theta LFP (5-11 Hz) and the amplitude of DG slow gamma ($SG_{LPP}$; left). The average $SG_{LPP}$ waveforms locked at $DS_L$ and $DS_M$ events (right) show that in time of maximal positive DS amplitude (T = 0 ms), $SG_{LPP}$ exhibits a negative polarity demonstrating robustness of separation of $SG_{LPP}$ and LFP waveforms associated with DS events. H) Comodulogram between the phase of *stratum pyramidale* theta LFP (5-11 Hz) and the amplitude of DG mid-frequency gamma ($MG_{MPP}$; left). Average $MG_{MPP}$ waveforms locked at $DS_L$ and $DS_M$ events (right) show that at the time of maximal positive DS amplitude (T = 0 ms), $MG_{MPP}$ exhibits negative polarity demonstrating robustness of separation of $MG_{MPP}$ and LFP waveforms associated with DS events. I) Comodulogram between the phase of *stratum pyramidale* theta LFP (5-11 Hz) and the amplitude of $FG_{MPP}$ IC, a third identified IC separate from $SG_{LPP}$ or $MG_{MPP}$ ICs. Average waveforms locked at $DS_L$ and $DS_M$ events (right) show that at the time of the maximal positive $DS_M$ amplitude (T = 0 ms), $FG_{MPP}$ IC exhibits a positive LFP polarity suggesting association between $FG_{MPP}$ IC and $DS_M$ events due to the same origin in MECII. Notice that $FG_{MPP}$ IC exhibits two peaks in the comodulogram (left), a high frequency (>100 Hz) component, and a low frequency component ~30 Hz, likely associated with spectral leakage from theta phase-locked $DS_M$ events (Figs. 3A, 3D). J) Comodulogram between the phase of *stratum pyramidale* theta LFP (5-11 Hz) and the amplitude of $FG_{MPP}$ (left), $FG_{MPP}$ during 100 ms windows centered on $DS_M$ (center) and $FG_{MPP}$ with times during $DS_M$ removed (right). Notice increased phase-amplitude coupling of the low frequency component ~30 Hz for $FG_{MPP}$ during $DS_M$ (middle), suggesting that the low-frequency component of $FG_{MPP}$ is likely associated with $DS_M$. K) Localization of DG ICs using DS CSD profiles. $DS_L$ (left) has CSD sinks in the outer molecular layers of DG at the LECII projection termination (gray arrowheads). $DS_M$ (middle) has CSD sinks in middle molecular layers of DG, at the MECII projection termination (red arrowheads). Medial perforant path stimulation (right) shows CSD sinks in the middle molecular layers of DG, at the MECII projection terminals (red arrowheads), same as $DS_M$. L) Example of CSDs of voltage loadings of 3 DG ICs in comparison to DS CSDs. M) Group data of distance between sink locations in CSDs of DG ICs relative to sink locations in $DS_L$ (left) and $DS_M$ (right). Distance normalized independently for each recording so the distance between $DS_L$ and $DS_M$ sink locations is unitary. Data combined from the superior and inferior blades of DG. Statistical comparisons computed for the difference from the $DS_L$ CSD sink location (left; t test: $SG_{LPP}$: $t_{10}$ = 0.91, p = 0.39; $MG_{MPP}$: $t_{10}$ = 2.49, p = 0.03; $FG_{MPP}$: $t_{11}$ = 6.05, p = $10^{-5}$) and DSM CSD sink location (right; t test: $SG_{LPP}$: $t_{10}$ = 5.12, p = $10^{-4}$; $MG_{MPP}$: $t_{10}$ = 0.59, p = 0.57; $FG_{MPP}$: $t_{11}$ = 3.07, p = 0.01). Notice that $SG_{LPP}$ localizes to $DS_L$ sink locations (LECII termination zone), $MG_{MPP}$ localizes to $DS_M$ sink locations (MECII termination zone) and $FG_{MPP}$ localizes between $DS_L$ and $DS_M$ sink locations, but closer to $DS_M$ sink locations.

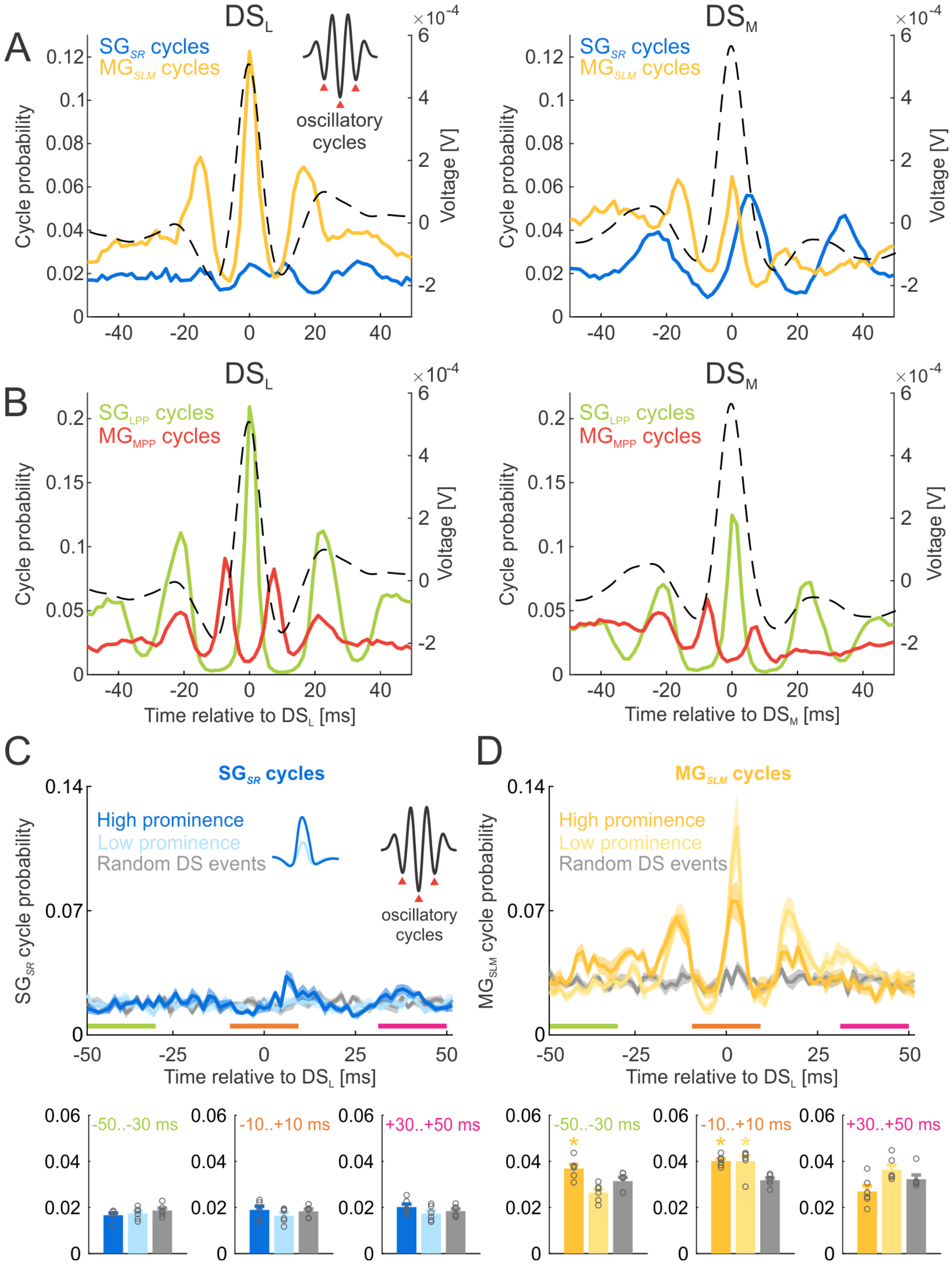


**Supplemental figure S3 related to Figure 4** A) Distribution of oscillatory cycles of $SG_{SR}$ (blue) and $MG_{SLM}$ (yellow) referenced to $DS_L$ (left) and $DS_M$ (right) obtained from LFPs recorded during home cage behavior. Compare robustness of relationships with results obtained from LFPs during active place avoidance (Fig. 4A). DS average waveform shown as dashed line for reference. Example oscillatory events with marked oscillatory cycles (red triangles) shown as inset. B) Distribution of oscillatory cycles of $SG_{LPP}$ (green) and $MG_{MPP}$ (red) referenced to $DS_L$ (left) and $DS_M$ (right) obtained from LFPs recorded during home cage behavior. DS waveform shown as dashed line for reference. C) Probability of $SG_{SR}$ oscillatory cycles (inset) around $DS_L$ with the 10% highest prominence (darker color) and 10% lowest prominence (lighter color).

Average probability of oscillatory cycles were assessed during -50..-30 ms before $DS_L$ (bottom, left), -10..+10 ms centered at $DS_L$ (bottom, middle) and +30..+50 ms after $DS_L$ (bottom, right). No differences between high and low prominence $DS_L$ events were observed. D) Probability of $MG_{SLM}$ oscillatory cycles around $DS_L$ with the 10% highest prominence (darker color) and 10% lowest prominence (lighter color). Average probability of oscillatory cycles were assessed during -50..-30 ms before $DS_L$ (bottom, left; $F_{2,17} = 12.16$; $p = 0.0007$; post-hoc tests: High > Low = Random), -10..+10 ms centered at $DS_L$ (bottom, middle; $F_{2,17} = 16.52$; $p = 0.0002$; post-hoc tests: High = Low > Random) and +30..+50 ms after $DS_L$ (bottom, right; $F_{2,17} = 5.74$; $p = 0.014$; High = Low = Random). Averages ± S.E.M. are plotted.

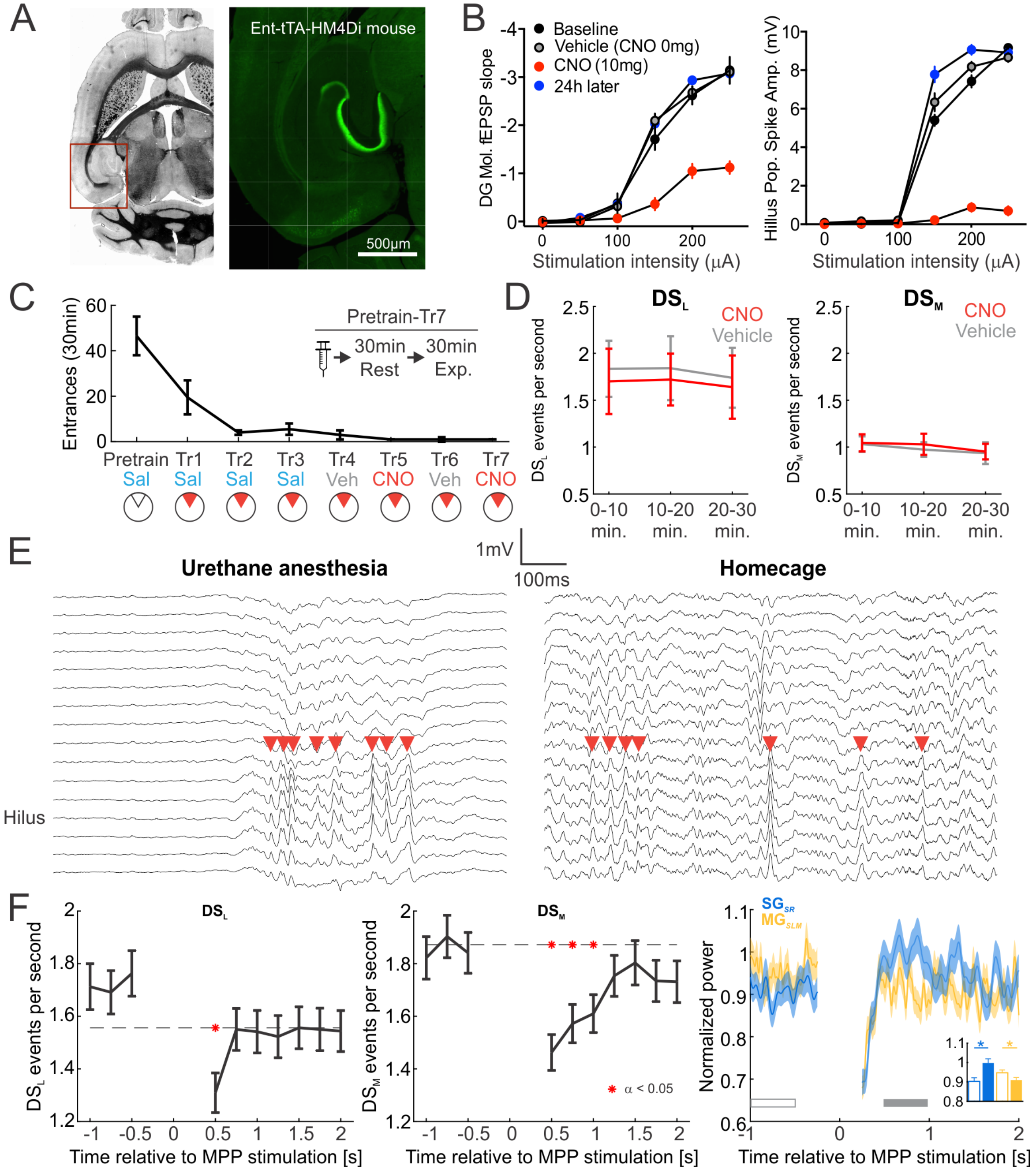

**Supplemental figure S4 related to Figure 4.** A) Brightfield and fluorescence images showing hM4Di expression in an Ent-tTA-hM4Di mouse, 1 week after doxycycline removal from the diet. This transgenic mouse expressed the inhibitory DREADD receptor hM4Di in the MEC layers 2 and 3 and was used to suppress neurons in the superficial layers of medial entorhinal cortex (MEC) that project to hippocampus through the MPP that is responsible for $DS_M$. Anti-hemagglutinin tag immunofluorescence indicating hM4Di expression is confined to MEC layer II and its afferent projections to DG and CA3. B) Input-output curves illustrating that CNO activation of hM4Di causes reversible suppression of perforant path input to dentate gyrus by hM4Di activation, measured as slope of the field excitatory postsynaptic potential (fEPSP) in molecular layer of DG (left) and population spike amplitude in hilus of DG (right). C) Experimental design and behavioral performance in active place avoidance task. To assure comparable behavioral expression, animals were trained in active place avoidance. Each session started with injection, followed by 30 min rest for drug to take effect and 30 min experiment. After pretraining (day 1, shock off, saline injection), animals were trained during 3 consecutive days (days 2-4, shock on, saline injection), followed by a session with vehicle injection (day 5), CNO 10 mg/kg injection (day 6), vehicle injection (day 7) and CNO 10 mg/kg injection (day 8). D) Rates of $DS_L$ (left) and $DS_M$ (right) during three 10-min segments of active place avoidance. CNO shown in red, vehicle shown in gray. Data from two vehicle sessions and two CNO sessions were averaged. These data suggest that DS generation is not strongly dependent on the MPP input. E) Dentate spikes (red triangles) under urethane anesthesia (left) and during home-cage behavior (right) demonstrating the ability to generate dentate spikes even under massive reduction of neural activity under urethane anesthesia. F) MPP stimulation (10 minutes off-on-off-on, 100 μs stimulus length, 15 s inter-stimulus interval, 200 μA stimulus intensity, while the mouse runs on a treadmill) reduces rates of $DS_L$ (left) 500 ms following the stimulation and reduces rates of $DS_M$ (middle) 500 – 1250 ms following the stimulation. Red asterisks mark significant deviations from the mean rate. $SG_{SR}$ power is increased and $MG_{SLM}$ power is decreased (right) post-stimulus compared to pre-stimulus (t test +500..+1000 ms vs -1000..-500 ms; $SG_{SR}$: $t_{640} = 3.82$, $p = 0.0001$; $MG_{SLM}$: $t_{640} = 2.35$, $p = 0.02$). These results demonstrate that electric stimulation of the medial perforant path creates a complex response, does not, simply generate artificial $DS_M$ events, and causes changes that would result in a higher probability of $SG_{dom}$.

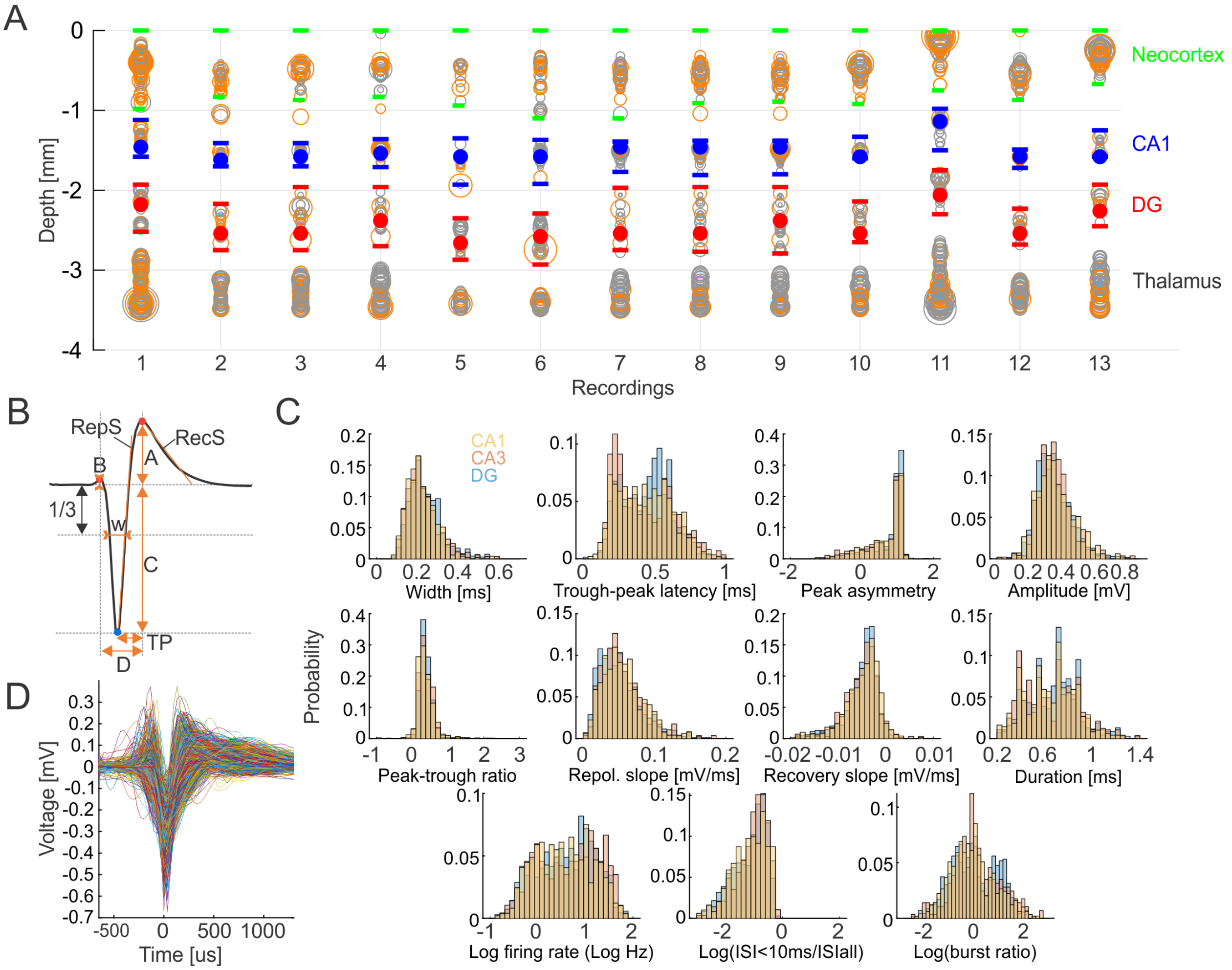


**Supplemental figure S5 related to Figure 6.** A) depth profile of detected single units in 13 consecutive recordings obtained using Neuropixels probes. Red dots mark depth of detected dentate spikes. Red horizontal lines mark putative DG boundary. Blue dots mark depth of detected sharp-wave ripples. Blue horizontal lines mark putative CA1 boundary. Green horizontal lines mark putative neocortical boundary. Open circles indicate single units, orange color represents units well isolated from background activity (< 20% estimated contamination rate with spikes from other neurons that were computed from the refractory period violations relative to expected), gray color represents multiunit activity, diameter corresponds to unit amplitude. B) Waveform features obtained from each isolated single unit that met the selection criteria. Width is measured as spike duration at 1/3 of spike amplitude from signal baseline (W); trough-peak latency is time between spike minima and the following maxima (TP); peak asymmetry is the ratio between the maxima preceding and following spike minima measured from signal baseline (b-a)/(b+a); spike amplitude is the sum of the two maxima (A+C); peak-trough ratio is the ratio of the two maxima (A/C); repolarization slope is measured by fitting a regression line to the first 165 μs from the spike minima towards the following maxima (RepS); recovery slope is measured by fitting a regression line to the first 660 μs from the following maxima (RecS); duration is the time between the maxima that precede and follow the peak (D). C) Histograms of extracted features from CA1 (yellow), CA3 (red) and DG (blue) hippocampus subfields. Notice the clear bimodal distributions in trough-peak latency and log(burst ratio) measures as well as the distinction between the features of DG and CA1+CA3 cells. D) Overlaid average waveforms of single units from CA1, CA3 and DG.

| | $DS_L$ | $DS_M$ |
|---|---|---|
| GC/E × GC/I | 0.65159 ± 1.0831<br>$t_{464}$ = -4.8941<br>P = 1.3649e-06 | 6.4937 ± 7.3429<br>$t_{464}$ = 11.239<br>P = 4.371e-26 |
| GC/E × MC/I | 0.37777 ± 0.51238<br>$t_{146}$ = -5.8711<br>P = 2.7995e-08 | 2.6215 ± 2.5673<br>$t_{146}$ = 6.1239<br>P = 8.0347e-09 |
| GC/E × MC/E | 0.36983 ± 0.56928<br>$t_{205}$ = -2.848<br>P = 0.0048481 | 2.507 ± 3.2264<br>$t_{205}$ = 5.9746<br>P = 1.0068e-08 |
| GC/E × CA3/E | 0.62202 ± 0.64689<br>$t_{54}$ = 0.36696<br>P = 0.71508 | 2.3396 ± 1.9221<br>$t_{54}$ = 2.117<br>P = 0.038886 |
| GC/E × CA3/I | 0.71057 ± 0.70674<br>$t_{72}$ = -2.0925<br>P = 0.039915 | 2.7956 ± 2.2425<br>$t_{72}$ = 5.6702<br>P = 2.7718e-07 |
| MC/E × GC/I | 0.83923 ± -1.0263<br>$t_{128}$ = -0.97357<br>P = 0.33211 | 4.6776 ± 4.9512<br>$t_{128}$ = 5.5327<br>P = 1.7059e-07 |
| MC/E × MC/I | 0.73545 ± 0.79413<br>$t_{113}$ = -1.0451<br>P = 0.2982 | 2.5696 ± 2.0566<br>$t_{113}$ = 5.7945<br>P = 6.2935e-08 |
| MC/I × GC/I | 0.83834 ± 1.1095<br>$t_{121}$ = -4.9891<br>P = 2.0546e-06 | 6.2324 ± 6.7617<br>$t_{121}$ = 8.897<br>P = 6.7828e-15 |
| CA3/E × CA1/E | 0.70343 ± 1.0584<br>$t_{129}$ = 2.05<br>P = 0.042394 | 0.74643 ± 0.7919<br>$t_{129}$ = 2.8737<br>P = 0.0047466 |
| CA3/E × CA3/I | 0.88858 ± 0.87775<br>$t_{518}$ = 2.265<br>P = 0.023926 | 2.3265 ± 2.9999<br>$t_{518}$ = 7.1865<br>P = 2.3315e-12 |
| CA3/E × CA1/I | 0.77526 ± 0.84038<br>$t_{224}$ = 0.7284<br>P = 0.46713 | 3.2339 ± 4.6558<br>$t_{224}$ = 3.4198<br>P = 0.00074447 |
| CA3/I × CA1/E | 0.8705 ± 0.92127<br>$t_{196}$ = -0.084586<br>P = 0.93268 | 1.8035 ± 2.1411<br>$t_{196}$ = 5.5339<br>P = 9.952e-08 |
| CA3/I × CA1/I | 1.2217 ± 1.3291<br>$t_{281}$ = 4.2895<br>P = 2.4654e-05 | 2.384 ± 2.6306<br>$t_{281}$ = 9.0801<br>P = 1.9735e-17 |
| CA1/E × CA1/I | 1.0562 ± 1.3194<br>$t_{361}$ = 2.832<br>P = 0.0048857 | 1.3288 ± 1.3742<br>$t_{361}$ = 5.4662<br>P = 8.5931e-08 |

**Table S1 related to Figure 6D.** Statistics of the co-occurrence probability between two types of cells in 6 ms windows around $DS_L$ and $DS_M$ events compared to random times. DS events that coincided with SWR events were excluded from the analysis. Each 3-row entry contains a mean ± S.D., and ratio of co-occurrence probability compared to random events ($\frac{P_{DS}(cell\ type\ 1, cell\ type\ 2)}{P_{RND}(cell\ type\ 1, cell\ type\ 2)}$; first row), t test statistics (second row) and associated p-value (third row). Significant comparisons were corrected using Bonferroni's method and are marked red ($\alpha$ = 0.0036).

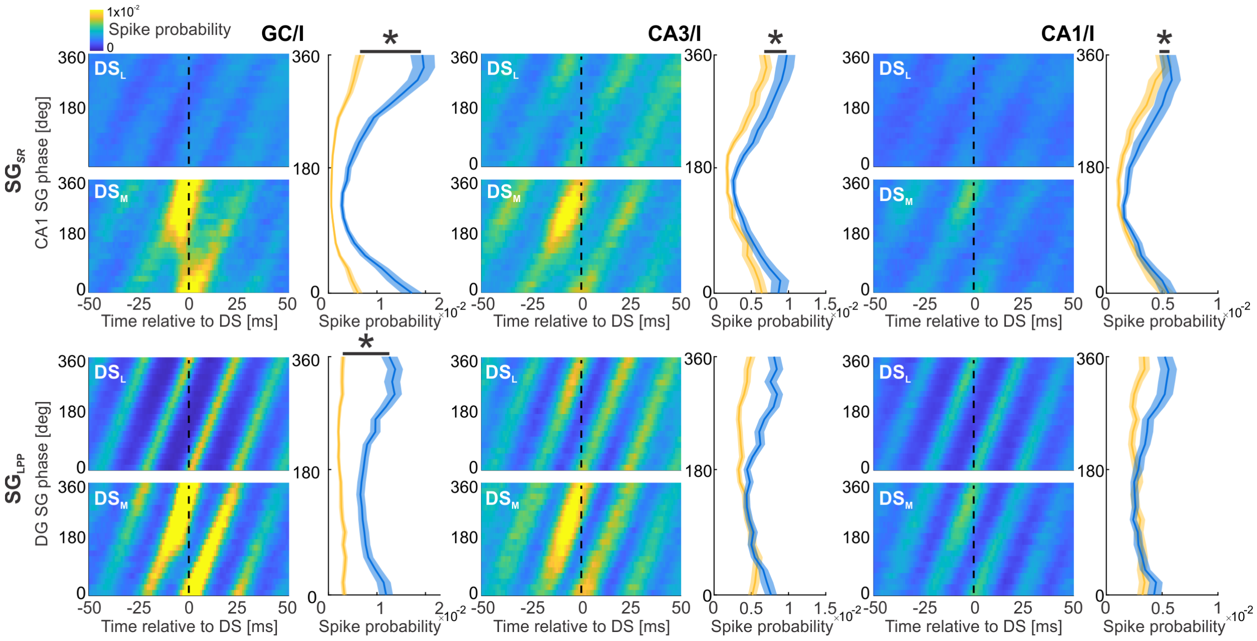


**Supplemental figure S6 related to Figure 7.** Discharge probability of narrow waveform interneurons localized to the vicinity of DG granule cells (left column), to CA3 (middle column) and to CA1 (right column) relative to the timing of $DS_L$ and $DS_M$ events (x axis) and phase of CA1 slow gamma ($SG_{SR}$; top row) and DG slow gamma ($SG_{LPP}$; bottom row). Dashed lines mark the peak amplitudes of DS events, where the average profiles were extracted for statistics (right side of each panel). Averages ± S.E.M. are plotted across all identified cells. Statistical comparisons are performed using Kuiper two-sample test, significant differences between $DS_M$ and $DS_L$ are marked with asterisks ($SG_{SR}$ x GC interneurons: $k = 1653$, $p < 0.001$; $SG_{SR}$ x CA3 interneurons: $k = 2838$, $p < 0.001$; $SG_{SR}$ x CA1 interneurons: $k = 2628$, $p < 0.001$; $SG_{LPP}$ x GC interneurons: $k = 1160$, $p = 0.02$; $SG_{LPP}$ x CA3 interneurons: $k = 732$, $p = 1$; $SG_{LPP}$ x CA1 interneurons: $k = 1596$, $p = 0.06$.
